# A TAL effector-like protein of symbiotic *Mycetohabitans* increases stress tolerance and alters the transcriptome of the fungal host *Rhizopus microsporus*

**DOI:** 10.1101/2020.03.04.968529

**Authors:** Morgan E. Carter, Sara C.D. Carpenter, Zoë E. Dubrow, Mark R. Sabol, Fabio C. Rinaldi, Olga A. Lastovestsky, Stephen J. Mondo, Teresa E. Pawlowska, Adam J. Bogdanove

## Abstract

Symbioses of bacteria with fungi have only recently been described and are poorly understood. In the symbiosis of *Mycetohabitans* (formerly *Burkholderia*) *rhizoxinica* with the fungus *Rhizopus microsporus*, bacterial type III (T3) secretion is known to be essential. Proteins resembling T3-secreted transcription activator-like (TAL) effectors of plant pathogenic bacteria are encoded in the three sequenced *Mycetohabitans spp.* genomes. TAL effectors nuclear localize in plants, where they bind and activate genes important in disease. The Burkholderia TAL-like (Btl) proteins bind DNA but lack the N- and C-terminal regions in which TAL effectors harbor their T3 and nuclear localization signals, and activation domain. We characterized a Btl protein, Btl19-13, and found that, despite the structural differences, it can be T3-secreted and can nuclear localize. A *btl19-13* gene knockout did not prevent the bacterium from infecting the fungus, but the fungus became less tolerant to cell membrane stress. Btl19-13 did not alter transcription in a plant-based reporter assay, but 15 *R. microsporus* genes were differentially expressed in comparisons both of the fungus infected with the wildtype bacterium vs the mutant and with the mutant vs. a complemented strain. Southern blotting revealed *btl* genes in 14 diverse *Mycetohabitans* isolates. However, banding patterns and available sequences suggest variation, and the *btl19-13* phenotype could not be rescued by a *btl* gene from a different strain. Our findings support the conclusion that Btl proteins are effectors that act on host DNA and play important but varied or possibly host-genotype-specific roles in the *M. rhizoxinica*-*R. microsporus* symbiosis.

## Introduction

Bacteria are critical partners in symbiotic interactions with a variety of animals and plants, ranging from bobtail squid to legumes. Many of these associations are well studied, and the reciprocal benefits and molecular interactions underlying them well understood. Recently, diverse bacterial-fungal symbioses have been described, with fungal hosts ranging from tree endophytes to human pathogens (Arora and Riyaz-Ul-Hassan, 2018). One such partnership is *Mycetohabitans rhizoxinica* (family *Burkholderiaceae*; formerly *Burkholderia rhizoxinica* (Estrada-de Los Santos et al., 2018)) and the mucoromycete *Rhizopus microsporus.* A rice pathogen and opportunistic human pathogen, *R. microsporus* can also be found as a free-living soil saprophyte and is used in soy fermentation. *M. rhizoxinica* produces an anti-mitotic toxin, rhizoxin, which is critical for the fungus to infect rice seedlings (Partida-Martinez and Hertweck, 2005).

Though challenging, *M. rhizoxinica* can be cultured and genetically manipulated independently of its fungal host, making it a useful model. *R. microsporus* isolates that harbor *M. rhizoxinica* or other *Mycetohabitans* spp. require their bacterial partner for asexual and sexual sporulation, while non-host isolates sporulate without bacteria (Partida-Martinez et al., 2007; Lackner et al., 2009; Mondo et al., 2017). As is true for many gram-negative bacterial pathogens and mutualists of plants and animals, *M. rhizoxinica* requires a type III secretion system (T3SS) to invade its host (He et al., 2004; Lackner et al., 2011a). T3SSs inject effector proteins directly into host cells to promote infection (Alfano and Collmer, 2004). Over the past few decades, the T3 effector repertoires of many plant and animal-associated bacteria have been identified, and their modes of action characterized (Hu et al., 2017).

Despite the demonstrated importance of the T3SS in the *Rhizopus-Mycetohabitans* partnership, no T3 effector proteins from *M. rhizoxinica* or any other endohyphal bacterium have been characterized (Nazir et al., 2017). However, homologs of transcription activator-like (TAL) effectors from plant pathogenic bacteria were identified in the first sequenced *M. rhizoxinica* genome (Lackner et al., 2011b; de Lange et al., 2014b; Juillerat et al., 2014; Estrada-de Los Santos et al., 2018). TAL effectors are T3-secreted, sequence-specific, DNA-binding proteins that act as transcription factors once inside plant cells, upregulating target genes. Often these targets are so-called susceptibility genes that contribute to bacterial proliferation, symptom development, or both; examples include sugar transporter, transcription factor, and other genes (Boch et al., 2014; Hutin et al., 2015). Some strains of the phytopathogenic *Ralstonia solanacearum* species complex also have TAL effectors (called ‘RipTALs’) (Heuer et al., 2007; Schandry et al., 2016). Outside of plant pathogens, TAL effector-like gene fragments have been found in the marine metagenome, although the organism or organisms of origin remain unknown (de Lange et al., 2015). The limited discovered distribution of this effector family makes its evolutionary origins and functional diversity a mystery.

TAL effectors have four structural components: an N-terminal T3S signal (Szurek et al., 2002), C-terminal nuclear localization signals (Van den Ackerveken et al., 1996; Szurek et al., 2001), a central, repetitive DNA recognition domain (Deng et al., 2012; Mak et al., 2012), and a C-terminal activation domain (Szurek et al., 2001). The DNA recognition domain consists of highly conserved repeats each with two variable amino acids, the repeat variable diresidue (RVD). The repeats each interact with single nucleotides, in a contiguous fashion, with specificity dictated by the RVDs. This one-to-one recognition “code” makes it possible to predict the DNA sequence(s) a TAL effector will bind given its RVD sequence (Boch et al., 2009; Moscou and Bogdanove, 2009). Also, TAL effectors can be synthesized to target specific DNA sequences by putting together repeats with the appropriate RVDs, making these proteins useful for biotechnology (Doyle et al., 2013; Boch et al., 2014; de Lange et al., 2014a).

The initial discovery of TAL effector homologs in *Mycetohabitans* (then *Burkholderia*) came just as TAL effectors were garnering interest as engineerable tools for DNA targeting applications such as genome editing. Though their repeats are less highly conserved, the *Burkholderia* proteins were found to bind DNA targets specifically, wrapping the DNA in a superhelical structure, like TAL effectors (Mak et al., 2012; de Lange et al., 2014b; Juillerat et al., 2014; Stella et al., 2014). In the biochemical studies, different names for the proteins were introduced, including BurrH, and the collective term Bat (Burkholderia TAL-like). Bat follows bacterial gene nomenclature convention but was in use prior to refer to bacteriopsin activator genes. Therefore, we refer to the proteins as Btl (for *Burkholderia* TAL-like) and encoding genes as *btl*. Though the studies to date established Btl proteins as sequence-specific DNA binding proteins with potential for synthetic biology, none has yet addressed the function(s) of these proteins within the bacterial-fungal symbiosis. The Btl protein structure is truncated at the N- and C-termini compared with canonical TAL effectors, and lacks a T3S signal, nuclear localization signals, and an activation domain (**Fig. 1A**). Thus, even whether Btl proteins act as translocated transcription factors is itself uncertain.

**Figure 1.**
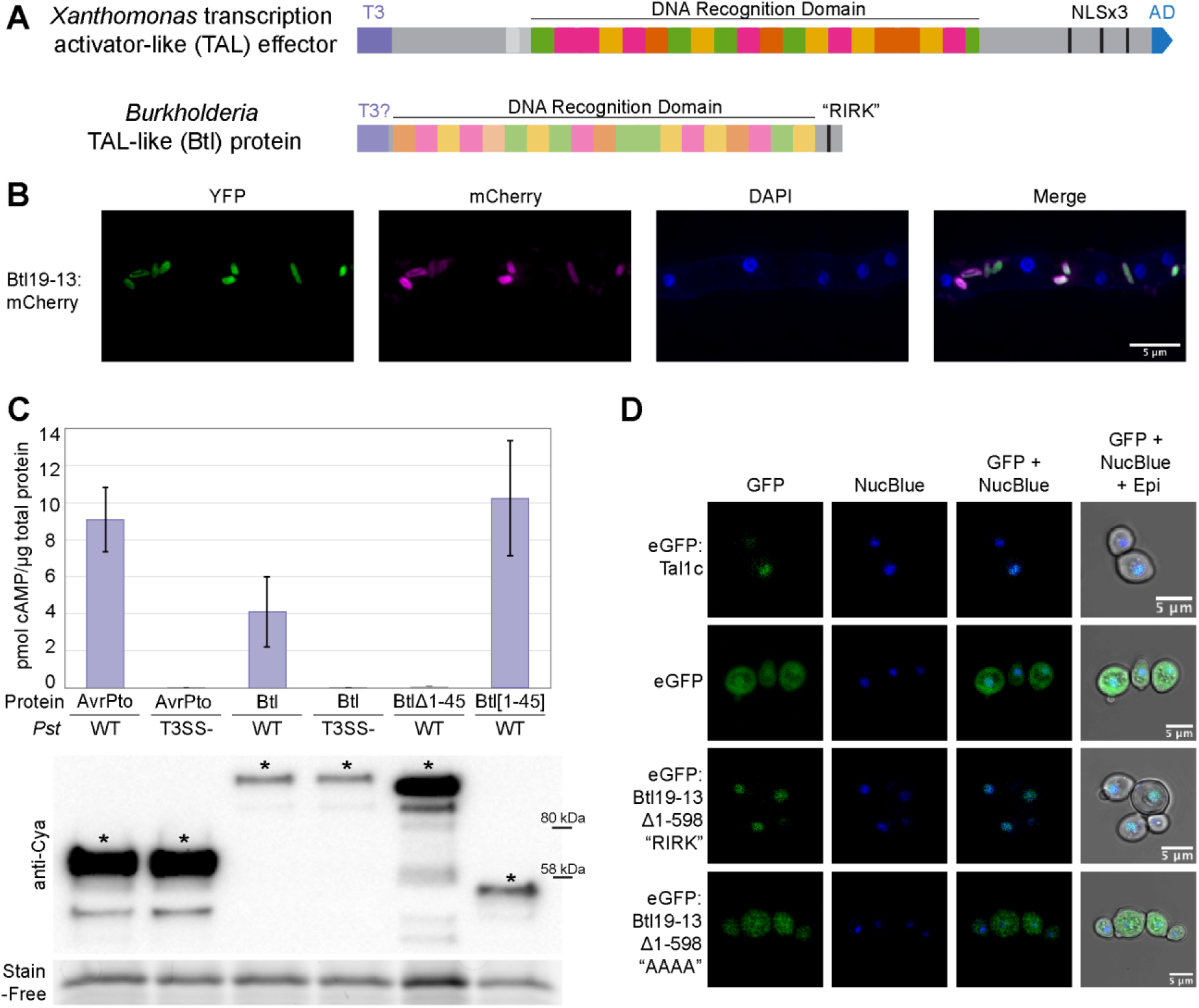
Structure, expression, T3 secretion, and nuclear localization of Btl19-13. **(A)** Domain structure of a *Xanthomonas* TAL effector and a Btl protein. T3, type III secretion signal; T3?, putative type III secretion signal; NLS, nuclear localization signal; “RIRK,” aa sequence of a putative NLS in Btl19-13; AD, activation domain. **(B)** Confocal microscopy of *Mycetohabitans* sp. B13 cells constitutively expressing EYFP and expressing Btl19-13:mCherry under the native *btl19-13* promoter, within a *R. microsporus* hypha. DAPI staining of the nuclei is included for reference. **(C)** Quantification of cAMP by ELISA in *Nicotiana benthamiana* leaf punches 6 hours after infiltration with *Pseudomonas syringae* pv. *tomato* DC3000 (*Pst*) expressing the following: AvrPto:Cya (61 kDa), Btl19-13:Cya (118.6 kDa), Btl19-13:Cya missing the first 45 aa (Btl19-13∆1-45, 110 kDa), and the first 45 aa of Btl19-13 fused to CyaA (Btl19-13[1-45], 51 kDa). T3SS-designates a *hrpQ-U* mutant incapable of Type III secretion (Badel et al., 2006). Each bar shows the results from three biological replicates with two technical replicates each. Error bars denote standard deviation. The experiment was repeated twice with similar results. Below, a western blot of leaf homogenates probed with anti-Cya, shown with a Stain-Free loading control (Bio-Rad). Asterisks (*) indicate bands corresponding to the expected size. **(D)** Confocal microscopy of *Saccharomyces cerevisiae* cells expressing the indicated proteins and stained with NucBlue to locate nuclei. Protein expression was induced with galactose 24 hours prior to imaging. “AAAA” indicates an alanine substitution of the “RIRK” motif in Btl19-13 (residues 692-695).

The identification of proteins homologous to TAL effectors in *Mycetohabitans* spp. presents an opportunity to probe the molecular basis of a fungal-bacterial symbiosis and to better understand the functional diversity of this intriguing family of proteins. In this study, we carried out several experiments to test the hypothesis that Btl proteins function like *Xanthomonas* TAL effectors, transiting the T3SS and altering host transcription. We also probed whether Btl proteins play any role in establishing the symbiosis or confer a benefit to the fungus. Finally, we explored the prevalence of *btl* genes across geographically diverse *Mycetohabitans* spp. and tested whether distinct *btl* genes found in different isolates might be functionally interchangeable.

## Results

### The *btl* gene in *Mycetohabitans* sp. B13 is expressed endohyphally

We identified *btl* genes in each of the three publicly available genome sequence assemblies of *Mycetohabitans* spp. that are complete or largely so: *Mycetohabitans rhizoxinica* HKI 0454/B1 endosymbiont of *R. microsporus* ATCC 62417 (NCBI accession: PRJEA51915; Lackner et al., 2011b), *Mycetohabitans* sp. B13 endosymbiont of *R. microsporus* ATCC 52813 (NCBI accession: PRJNA303198), and *Mycetohabitans* sp. B14 endosymbiont of *R. microsporus* ATCC 52814 (NCBI accession: PRJNA303197). Strain B13 has one intact *btl* gene and a small *btl* gene fragment, whereas strains B1 and B14 each have three *btl* genes (**Table S1**). We named each using the number of repeats in the recognition domain followed by the strain number; for example, the single intact *btl* gene of strain B13, which has 19 repeats, was named *btl19-13*. To facilitate functional characterization, we chose to avoid the possible redundancy in strains B1 and B14 and focus instead on the single *btl* gene in strain B13.

First, to determine whether *btl19-13* is expressed, we cloned it on a plasmid under the control of its native promoter and with a C-terminal translational fusion to mCherry. After transforming B13 with this plasmid, we introduced the transformant into the corresponding, cured *R. microsporus* isolate, and assayed for mCherry fluorescence by confocal microscopy. The plasmid we used also harbors a constitutively expressed EYFP gene, to enable independent visualization of the bacterial cells. Btl19-13:mCherry and EYFP signals were evident and coincident within the fungal hyphae (**Fig. 1B**), indicating that the bacterium expresses *btl19-13* during the symbiosis.

### Btl19-13 transits the T3SS of *Pseudomonas syringae*

Our next question was whether Btl19-13 could, like TAL effectors, transit a T3SS and thereby potentially function within the fungal host. In the fluorescence microscopy described above, Btl19-13:mCherry was detectable only in the bacterium, but the fluorophore may have prevented secretion. Computational analysis of the Btl19-13 protein sequence using EffectiveT3 2.0.1 (Eichinger et al., 2015) detected with low confidence the presence a T3S signal in the N-terminus. Upstream of the *btl19-13* gene we discovered a sequence that aligns with the plant-inducible promoter (PIP) element found in promoters of genes co-regulated with the T3SS in *Xanthomonas* and *Ralstonia* spp. (Fenselau and Bonas, 1995; Cunnac et al., 2004). The sequence upstream of the *btl19-13* gene, as well as *btl* genes in B1 and B14, matches the consensus targeted by the regulator HrpB, TTCG-N16-TTCG (**Table S2**) (Lackner et al., 2011a).

We therefore took advantage of an established assay for T3S using the plant pathogen *Pseudomonas syringae* pv. *tomato* DC3000 (*Pst*) inoculated to leaves of the model plant *Nicotiana benthamiana* (Schechter et al., 2004) to test whether Btl19-13 is a T3S substrate. This assay detects activity of an adenylate cyclase (Cya; the catalytic domain of the *Bordetella pertussis* toxin CyaA) fused to the test protein (Ladant and Ullmann, 1999). Cya activity (conversion of ATP to cAMP) requires calmodulin and therefore acts as a reporter of localization into the host cell following T3S. We used the *Pst* effector AvrPto fused to Cya as a positive control, and included a *Pst* T3SS-deficient strain to assess T3SS-dependence of activity (Schechter et al., 2004; Badel et al., 2006). Following infiltration, Btl19-13:Cya resulted in increased cAMP when expressed in *Pst* but not *Pst* T3SS-, indicating that Btl19-13 is a T3SS substrate (**Fig. 1C**). Furthermore, expression of truncated constructs revealed that the first 45 amino acids of Btl19-13 are necessary and sufficient for T3S (**Fig. 1C**). Together, the results presented here support the hypothesis that Btl proteins are T3 effectors.

### Btl19-13 has a functional nuclear localization signal

While Btl proteins bind DNA, suggesting they localize to the nucleus, their short C-terminus does not contain any predicted NLSs. However, we identified a short, NLS-like sequence, “RIRK,” in the C-terminal region of Btl19-13 (residues 692-695) that is also present in the other Btl proteins (**Table S1**). Because there is not yet a method to genetically manipulate *R. microsporus*, in order to assess Btl19-13 subcellular localization, we transformed the yeast *Saccharomyces cerevisiae* with inducible expression plasmids encoding eGFP fusion proteins. When viewed with confocal microscopy, yeast cells induced to produce free eGFP showed diffuse localization of the protein. Conversely, as expected, eGFP-tagged TAL effector Tal1c cloned from *Xanthomonas oryzae* pv. *oryzicola* localized to the nucleus (**Fig. 1D**). However, we were unable to see fluorescence in induced cells transformed with eGFP:Btl19-13, suggesting that the full-length Btl19-13 protein is toxic to yeast cells. We generated an N-terminally truncated derivative, Btl19-13∆1-598, expressing only a little over two repeats of the DNA recognition domain and the entire C-terminal region, containing the “RIRK” motif. This construct fused to eGFP localized to the nuclei of the yeast cells, demonstrating that Btl19-13 contains at least one functional NLS (**Fig. 1D**). Localization to the nucleus was abolished when the “RIRK” motif was mutated to alanines (AAAA). This same substitution in the full-length Btl19-13 protein fused to eGFP resulted in diffuse signal in the yeast cells (**Fig. S1**). The presence of a functional NLS in Btl19-13, along with the observation that wildtype Btl19-13 appears to be toxic to yeast while a non-nuclear localizing derivative is not, supports the hypothesis that Btl proteins function in the host nucleus.

### Btl19-13 does not affect activity of a reporter gene in an *in planta* transient expression assay

To address whether Btl proteins might directly alter transcription of host genes, as TAL effectors do, we used an *Agrobacterium*-mediated transient expression assay in *N. benthamiana.* To first test if Btl19-13 could upregulate expression of a GUS reporter gene driven by a minimal promoter from the pepper *Bs3* gene (Römer et al., 2010; Cernadas et al., 2014), we amended the promoter with a binding element (BE) for the Btl19-13 (**Fig. 2A**). We designed the BE based on the Btl19-13 RVD sequence and showed by electrophoretic mobility shift assay that Btl19-13 binds it specifically (**Fig. S2**). As positive and negative controls, respectively, we also tested a designer TAL effector, dT19-13, constructed to target the Btl19-13 BE (**Fig. 2B**), and Tal1c, which has no predicted BE in the promoter of the reporter construct. For reference, we included AvrBs3, the native TAL effector of *Xanthomonas euvesicatoria* that activates the minimal *Bs3* promoter via a separate BE. Each of these proteins was also tested on a construct harboring a scrambled version of the Btl19-13 BE (sBE; **Fig. 2B**). Unlike dT19-13, which specifically activated the reporter harboring the Btl19-13 BE, Btl19-13 did not alter the reporter activity of either construct (**Fig. 2B**). We next asked whether Btl19-13 could repress transcription by competing with an activator that binds the same target sequence or one nearby. In the *N. benthamiana* GUS assay, using the construct harboring the Btl19-13 BE (and the AvrBs3 BE just downstream), we co-introduced an activator, dT19-13 or AvrBs3, with Btl19-13 or, as the control, with Tal1c. Btl19-13 did not significantly reduce GUS activity relative to Tal1c in either combination (**Fig. 2C**).

**Figure 2.**
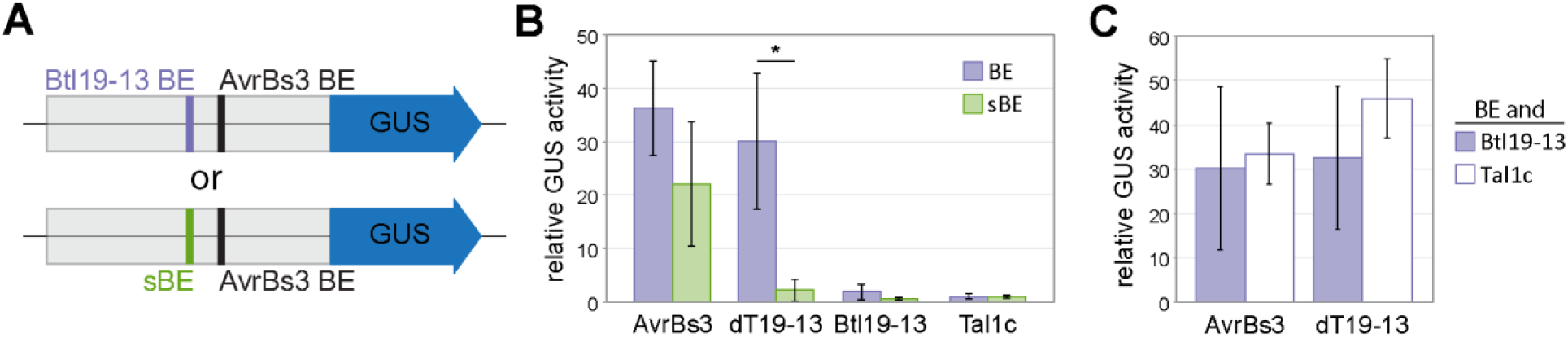
Effect of Btl19-13 on reporter gene activity in a transient expression assay in *N. benthamiana* leaves. **(A)** Schematic of the pepper *Bs3* minimal promoter-driven GUS construct used, harboring the AvrBs3 binding element (BE) and either a Btl19-13 BE or a scrambled version (sBE). **(B-C)** Fluorometric assays of GUS activity *in planta. N. benthamiana* leaves were co-infiltrated with *Agrobacterium tumefaciens* strains delivering a promoter:GUS fusion and one **(B)** or more **(C)** effector protein expression constructs, as indicated, and assayed 48 hours later. Activity is shown relative to just Tal1c and the corresponding reporter construct. Each value is an average of three replicates, and the experiment was repeated twice with similar results. An asterisk (*) over two values indicates a significant difference (paired student’s t test, *p*<0.05). Error bars denote standard deviation.

### Btl19-13 contributes to *R. microsporus* tolerance to cell membrane stress

Having established in heterologous contexts that Btl19-13 is likely an effector that acts in the *R. microsporus* nucleus, but having not detected any transcription factor activity in the reporter assay, we decided to knock out the *btl19-13* gene in strain B13 to directly assess its impact on the symbiosis. We generated a *btl19-13* knockout derivative, B13Δ*btl19-13* (**Fig. S4A**). B13Δ*btl19-13* was no different from wild type in its ability to infect the fungus. Furthermore, *R. microsporus* infected with B13Δ*btl19-13* showed no difference in asexual sporulation, could still mate to produce sexual spores, and grew normally on nutrient-rich and poor media (**Fig. S4B-E**).

Since the *btl19-13* knockout had no gross effect on establishment and maintenance of the host-symbiont interaction, we next explored the hypothesis that Btl19-13 contributes to the symbiosis by increasing tolerance of the fungus to environmental stresses. We measured growth on half-strength potato dextrose agar (½ PDA) amended as follows to test specific stresses: with hydrogen peroxide (H_2_O_2_) for oxidative stress, with sodium chloride (NaCl) for osmotic stress, and with sodium dodecyl sulfate (SDS) for cell membrane stress. *R. microsporus* infected with B13Δ*btl19-13* grew as well as the fungus infected with wildtype B13 on 500 mM NaCl and on 3 mM H_2_O_2_ (**Fig. S4F-G**), but more slowly on media amended with 0.005% SDS (**Fig. 3A-B**). The reduced growth in the presence of the detergent was restored to wild type levels when B13Δ*btl19-13* was transformed with a plasmid (pBtl19-13) carrying the cloned *btl19-13 gene* with its native promoter (620 bp), prior to reinfection of the fungus.

**Figure 3.**
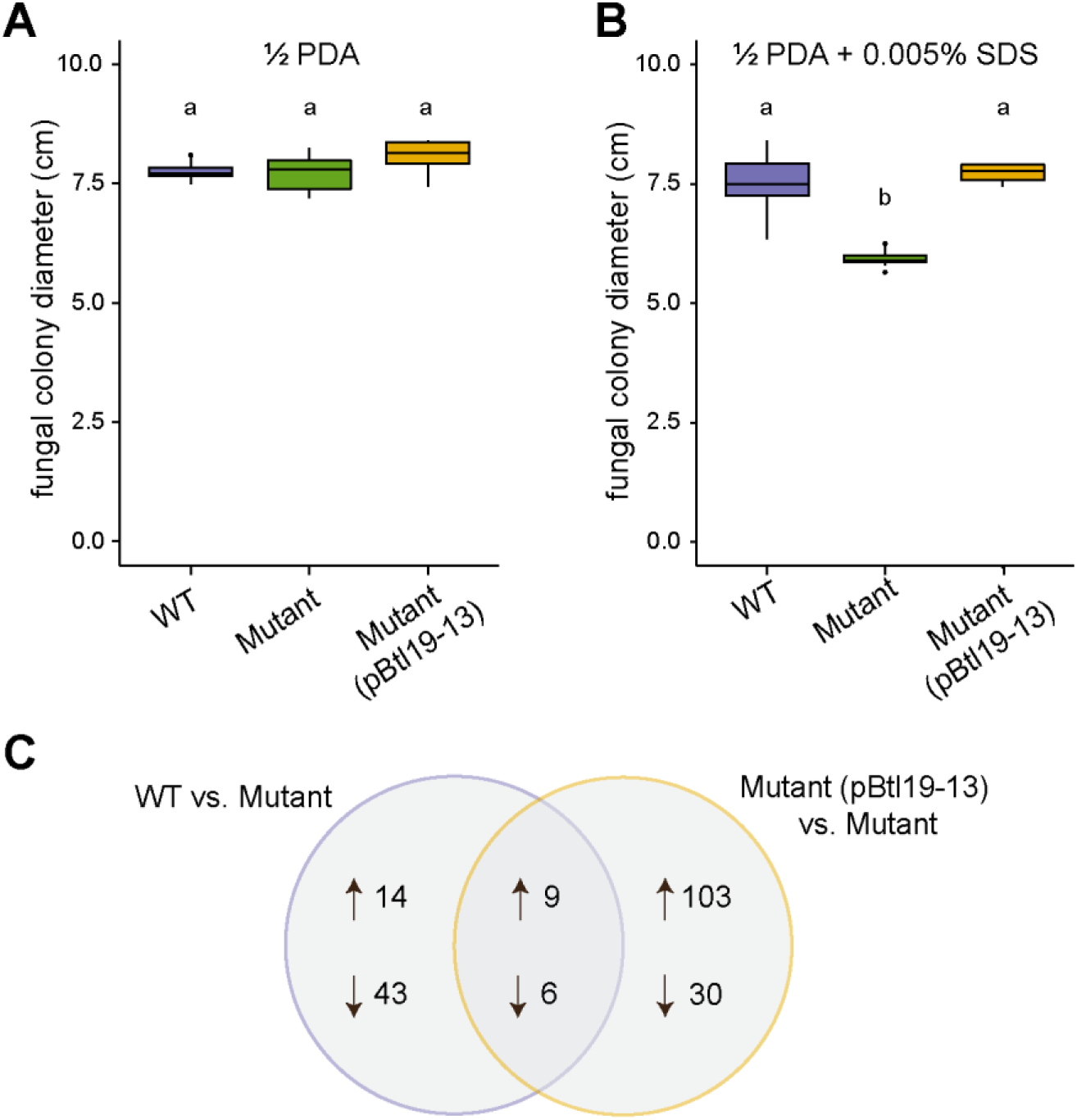
Effect of Btl19-13 on *R. microsporus* cell membrane stress tolerance and global gene expression. **(A-B)** Colony diameter of *R. microsporus* infected with wildtype *Mycetohabitans* sp. B13 (WT), B13Δ*btl19-13* (Mutant), or the mutant strain carrying *btl19-13* on a plasmid (pBtl19-13). Fungi were grown at 28°C on half-strength potato dextrose agar without (**A**) or with (**B**) 0.005% sodium dodecyl sulfate (SDS), for 3 and 6 days, respectively. Data shown represent 10 biological replicates for each bacterial genotype and were analyzed by ANOVA with a post-hoc Tukey’s test to determine significance as indicated by lower-case letters (*p*<0.001). The experiment was repeated twice and yielded similar results. **(C)** Venn diagram of the genes that were differentially expressed between *R. microsporus* infected with wildtype B13 or the complement, B13Δ*btl19-13*(pBtl19-13), and the mutant B13Δ*btl19-13.* Differential expression was determined by RNA sequencing and data analyzed with DESeq2 (Love et al., 2014) using an adjusted *p*-value < 0.05 as the threshold for significance. Three biological replicates were sequenced, each containing tissue from three plates.

### Btl19-13 alters the *R. microsporus* transcriptome

To probe the mechanism underlying the contribution of Btl19-13 to cell membrane stress tolerance of *R. microsporus*, we conducted RNAseq on mycelia infected with B13, B13Δ*btl19-13,* or the complemented B13Δ*btl19-13* strain and carried out pairwise comparisons. Fifteen genes were similarly differentially expressed (DE) in the comparison between B13 and B13Δ*btl19-13* and in the comparison between the complemented strain and B13Δ*btl19-13* (**Fig. 3C, Table S3**). Fourteen of the fifteen have predicted Btl19-13 BEs within their promoters (**Table S3**). However, annotations and pfam analysis did not provide obvious clues to the mechanism by which the Btl19-13 contributes to SDS tolerance; all but two of the DE genes encode hypothetical proteins or domains of unknown function (**Table S3**).

### *btl* genes are present across *Mycetohabitans* spp. but can differ functionally

Given that Btl19-13 does not appear to be integral to the formation or maintenance of the symbiosis, but instead appears to enhance the tolerance of the host to cell membrane stress, it was unclear whether all *Mycetohabitans* spp. could be expected to have *btl* genes serving the same function. The *btl* genes of strains B1, B3, and B14 encode distinct arrays of RVDs, which would indicate differences in their targeted BEs (**Table S1**). To survey *btl* genes across *Mycetohabitans* isolates, we assayed by Southern blot a collection of 14 geographically diverse strains isolated from across the variety of niches that *R. microsporus* inhabits (**Fig. 4A** and **Table S4**), including B1, B13, and B14. We used the cloned *btl19-13* gene of B13 as a probe. The numbers and sizes of bands detected for B1, B13, and B14 matched what was predicted based on the genome sequences (**Fig. 4B**). While B13 only has one predicted functional *btl* gene, there is also a small *btl* gene fragment (435 nucleotides) that we do not believe encodes a functional Btl protein based on its small size, lack of non-repeat domains, and dissimilar promoter with no predicted HrpB binding element. Each of the additional 11 strains yielded at least one strongly hybridizing band in the Southern blot, except NRRL 5549, which showed a faint, high-molecular weight band that is clearer at longer exposures (**Fig. 4B** and **Fig. S5**). A few bands were common to multiple strains, but no two strains shared the same overall banding pattern, suggesting that while *btl* genes are widely distributed in *Mycetohabitans*, they or their genomic contexts are variable.

**Figure 4.**
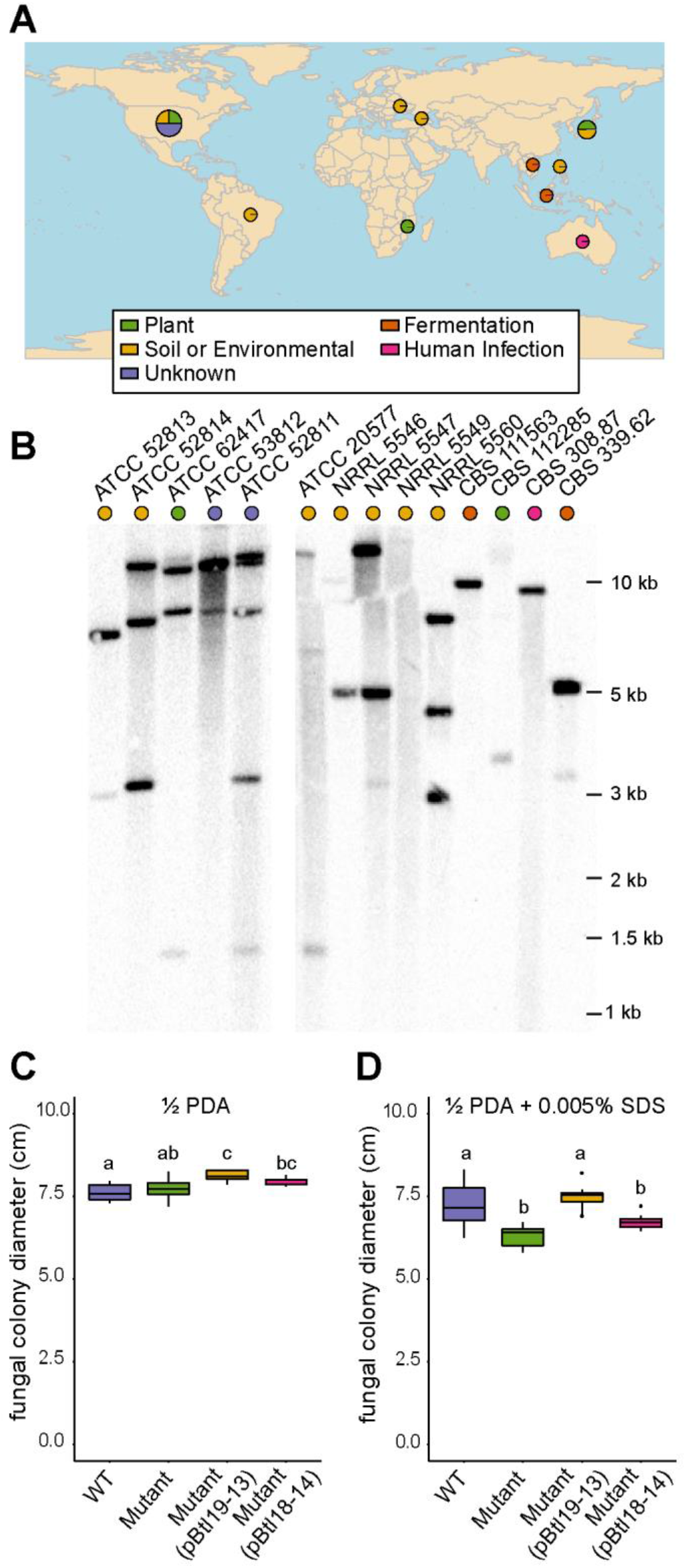
Presence of *btl* genes in geographically diverse isolates of *Mycetohabitans* spp. and the effect of substituting *btl18-14* for *btl19-13.* **(A)** World map showing the location and substrate from which the *R. microsporus* hosts of the *Mycetohabitans* spp. assessed were isolated, generated using rworldmap (South, 2011). More information on the strains is available in Table S3. **(B)** Southern blots of genomic DNA from each strain, digested with AatII and probed with *btl19-13* amplified from B13. Strains are identified by the culture collection accession number of their fungal hosts. ATCC 52813 represents B13, ATCC 52814 B14, and ATCC 62417 B1. **(C-D)** Growth of *R. microsporus* infected with wildtype B13 (WT), B13Δ*btl19-13* (Mutant), the mutant strain with pBtl19-13, or the mutant strain with pBtl18-14, on half-strength potato dextrose agar without **(C)** or with **(D)**0.005% sodium dodecyl sulfate (SDS) after 3 or 6 days, respectively, at 28°C. Data shown represent values from 10 replicate plates each and were analyzed by ANOVA with a post-hoc Tukey’s test. The experiment was repeated twice and yielded the same result. In each plot, different lower-case letters above any two groups indicate a significant difference between the means (*p*<0.001).

The apparent diversity of *btl* genes across strains suggests that Btl proteins have differentiated in function. To explore this possibility more directly, we cloned the 18-repeat *btl* gene from strain B14 (*btl18-14*), which is the sequenced *btl* gene encoding the RVD sequence most similar to that of Btl19-13 (**Table S1**), and tested whether it would rescue the B13Δ*btl19-13* phenotype of reduced tolerance to SDS. It did not (**Fig. 4C-D**), indicating that at least between these two strains, Btl proteins are not interchangeable.

## Discussion

The T3 translocation of Btl19-13, its nuclear localization, its contribution to host cell membrane stress, and the fact that it alters the host transcriptome support the hypothesis that Btl proteins act as effectors in *Mycetohabitans*-*Rhizopus* symbioses. Btl19-13 displays these properties despite largely truncated N- and C-terminal regions and degenerate or cryptic nuclear localization and T3S signals compared to *Xanthomonas* and *Ralstonia* TAL effectors. Endosymbiotic bacteria are observed to undergo genome reduction and to accumulate deleterious mutations relative to free-living bacteria due to their smaller population sizes, reduced recombination, and stable environment, which imposes purifying selection on a smaller number of genes (Muller, 1964; Moran, 1996). It seems plausible that Btl proteins represent degenerate descendants of ancestral TAL effector-like proteins. Indeed, it is striking, given the structural differences, that Btl19-13 retains the ability to be T3 secreted and nuclear localize. Likewise striking, Btl proteins specifically bind DNA despite the degeneracy of their central repeat sequences relative to TAL effectors. Together these observations are consistent with Btl proteins being under relatively strong selection as effectors that act on host DNA. The properties of Btl proteins contrast with those of a distinct class of structurally degenerate TAL effector-like proteins, TruncTALEs, which are found in some strains of the rice pathogen *Xanthomonas oryzae*. TruncTALEs lack the ability to bind DNA but block the function of a certain class of plant disease resistance gene, possibly through protein-protein interaction (Read et al., 2016).

While the data presented demonstrate that Btl19-13 alters the host transcriptome, the mechanism remains unclear, as we could not detect an effect of Btl19-13 on transcription in an *in planta* reporter gene assay. Specifically, Btl19-13 failed to activate a reporter with a Btl19-13 BE, while a dTALE matching that BE did. Furthermore, Btl19-13 did not measurably reduce dTALE-driven expression by direct competition, nor did it actively repress expression driven by a TAL effector targeting a downstream BE. Given the coevolution of the endosymbiont and host, Btl proteins may be adapted to function as activators or repressors by partnering with specific *Rhizopus* proteins or other effectors not present in the *in planta* assay. Alternatively, Btl protein repressor activity might depend on the specific promoter context, as Btl proteins have been observed to repress transcription of a reporter in *E. coli* when the promoter is amended with a corresponding BE (de Lange et al., 2015).

The mechanism by which Btl19-13 increases the tolerance of *Rhizopus* to detergent is likewise unclear, but could be related to major changes in *Rhizopus* lipid metabolism observed during infection (Lastovetsky et al., 2016). Reprogramming of lipid metabolism in the fungus has been suggested to help meet the bacterium’s nutritional requirement or favor invasion by the bacterium (Lastovetsky et al., 2016). While our study showed that Btl19-13 is not required for host infection, the protein could confer a quantitative benefit we did not measure. Annotated lipid metabolism genes were not present among the genes differentially expressed in response to Btl19-13, but some of the DE genes might still influence lipid metabolic pathways. It is also possible that the effect of Btl19-13 is highly localized, and not adequately captured by RNAseq of whole fungal colonies. Sampling of nuclei adjacent to bacterial cells could overcome this limitation. Furthermore, metabolome or lipidome analysis could reveal changes dependent on Btl19-13 and suggest genomic targets to investigate. Though beyond the scope of the current study, such approaches are an attractive first step toward understanding the connection between the observed influence of Btl19-13 on fungal SDS tolerance and the relevant biological function of Btl19-13. In other fungi, intolerance to SDS has been associated with changes not only in lipid metabolism (Gsell et al., 2015), but also fungal virulence (Lopez-Fernandez et al., 2018) and amino acid metabolism (Schroeder and Ikui, 2019). Thus, Btl19-13 could confer a variety of biologically relevant benefits to the fungus, the bacterium, or both.

The hypothesis that Btl proteins are important for the symbiosis is strengthened by the observation that, despite the reduced genomes of *Mycetohabitans* spp.*, btl* genes are present in all strains we examined in a collection of *R. microsporus* isolates from diverse geographic locations and substrates. The structural features of Btl19-13 are likely conserved in the other Btl proteins, based on similarity of the sequenced *btl* genes. However, there are a number of lines of evidence that these proteins do not all have the same functional target: (1) the number of *btl* genes in each strain varies, (2) the RVD sequences of the sequenced *btl* genes differ, and (3) B13Δ*btl19-13* could not be rescued with *btl18-14*. Btl proteins in different strains may target host genes with distinct functions or different genes with related functions. Or, they may target regions of the same gene(s) that are polymorphic across *Rhizopus* isolates. In the latter scenario, though Btl19-13 is not required for infection and is thus not a host specificity determinant in the strict sense, Btl proteins may influence patterns of association between specific bacterial and fungal isolates by contributing differentially to the persistence of the symbiosis depending on the host genotype. Further characterization of the sequence diversity of *btl* genes, comparative genomics of the corresponding host fungal isolates, and analysis of the Btl protein-dependent changes in the host transcriptome across a wide array of isolates will be important future steps toward determining the role(s) Btl proteins and their targets play in bacterial-fungal symbioses.

## Methods

Bacteria were isolated from fungi by tissue disruption and filtration, and fungi were cured with antibiotics and reinfected by cocultivation, as described (Lastovetsky et al., 2016). A variety of plasmids (**Table S5**) with *btl* gene derivatives were constructed by PCR, restriction enzyme digests, and mutagenesis. Microscopy of *R. microsporus* was done with a DeltaVision imaging station and yeast microscopy was done with a Zeiss 710 microscope. Type III Secretion was investigated using a Cya assay, as described (Chakravarthy et al., 2017). The reporter gene assay was done as described (Cernadas et al., 2014). For RNA-seq, RNA was extracted from day-old cultures of *R. microsporus* using an RNeasy Plant Mini Kit (Qiagen) and sent to Novogene for library preparation and Illumina sequencing. For amended media experiments, plates of *R. microsporus* were started with 2 mm x 2 mm plugs of 2-day-old fungus and the colony diameter was measured daily. For the Southern blot, bacterial genomic DNA was prepared using the MasterPure™ Gram Positive DNA Purification Kit (Lucigen) and fragmented by AatII digestion and the blot was probed with *btl19-13*. Full details of methods and materials are presented in the supplementary material.

## Acknowledgements

The authors thank Megan Feely, Pallavi Singh, and Ashli Gerschutz for laboratory assistance; Melanie Filiatrault for *N. benthamiana* plants; Alan Collmer and Suma Chakravarthy for *Pst* strains and plasmids, as well as guidance on the Cya assay; Wojtek Pawlowski for the use of the DeltaVision imaging workstation; Kathie Hodge for the use of the Olympus SZX12 dissecting microscope; Bryan Swingle for use of the plate reader; and the Cornell University Biotechnology Resource Center for the shared Zeiss LSM 710 Confocal Microscope (NIH 1S10RR025502). This work was supported by the USDA National Institute of Food and Agriculture (Predoctoral Fellowship award no. 2018-67011-28015 to M.E.C.), the National Science Foundation (grant no. IOS-1261004 to T.E.P.), and the National Institutes of Health (grant no. R01GM098861 to A.J.B.).

## References

Alfano, J.R., and Collmer, A. 2004. Type III Secretion System Effector Proteins: Double Agents in Bacterial Disease and Plant Defense. Annual Review of Phytopathology 42:385–414.

Arora, P., and Riyaz-Ul-Hassan, S. 2018. Endohyphal bacteria; the prokaryotic modulators of host fungal biology. Fungal Biology Reviews.

Badel, J.L., Shimizu, R., Oh, H.S., and Collmer, A. 2006. A *Pseudomonas syringae* pv. *tomato* avrE1/hopM1 mutant is severely reduced in growth and lesion formation in tomato. Mol Plant Microbe Interact 19:99–111.

Boch, J., Bonas, U., and Lahaye, T. 2014. TAL effectors – pathogen strategies and plant resistance engineering. New Phytologist 204:823–832.

Boch, J., Scholze, H., Schornack, S., Landgraf, A., Hahn, S., Kay, S., Lahaye, T., Nickstadt, A., and Bonas, U. 2009. Breaking the code of DNA binding specificity of TAL-type III effectors. Science 326:1509–1512.

Cernadas, R.A., Doyle, E.L., Niño-Liu, D.O., Wilkins, K.E., Bancroft, T., Wang, L., Schmidt, C.L., Caldo, R., Yang, B., White, F.F., Nettleton, D., Wise, R.P., and Bogdanove, A.J. 2014. Code-Assisted Discovery of TAL Effector Targets in Bacterial Leaf Streak of Rice Reveals Contrast with Bacterial Blight and a Novel Susceptibility Gene. PLOS Pathogens 10:e1003972.

Chakravarthy, S., Huot, B., and Kvitko, B.H. 2017. Effector Translocation: Cya Reporter Assay. Pages 473–487 in: Bacterial Protein Secretion Systems: Methods and Protocols, L. Journet and E. Cascales, eds. Springer New York, New York, NY.

Cuculis, L., Abil, Z., Zhao, H., and Schroeder, C.M. 2015. Direct observation of TALE protein dynamics reveals a two-state search mechanism. Nat Commun 6:7277.

Cunnac, S., Boucher, C., and Genin, S. 2004. Characterization of the *cis*-Acting Regulatory Element Controlling HrpB-Mediated Activation of the Type III Secretion System and Effector Genes in *Ralstonia solanacearum*. Journal of Bacteriology 186:2309.

de Lange, O., Binder, A., and Lahaye, T. 2014a. From dead leaf, to new life: TAL effectors as tools for synthetic biology. The Plant Journal 78:753–771.

de Lange, O., Wolf, C., Dietze, J., Elsaesser, J., Morbitzer, R., and Lahaye, T. 2014b. Programmable DNA-binding proteins from *Burkholderia* provide a fresh perspective on the TALE-like repeat domain. Nucleic Acids Res 42:7436–7449.

de Lange, O., Wolf, C., Thiel, P., Kruger, J., Kleusch, C., Kohlbacher, O., and Lahaye, T. 2015. DNA-binding proteins from marine bacteria expand the known sequence diversity of TALE-like repeats. Nucleic Acids Res.

Deng, D., Yan, C., Pan, X., Mahfouz, M., Wang, J., Zhu, J.K., Shi, Y., and Yan, N. 2012. Structural basis for sequence-specific recognition of DNA by TAL effectors. Science 335:720–723.

Doyle, E.L., Stoddard, B.L., Voytas, D.F., and Bogdanove, A.J. 2013. TAL effectors: highly adaptable phytobacterial virulence factors and readily engineered DNA-targeting proteins. Trends in Cell Biology 23:390–398.

Eichinger, V., Nussbaumer, T., Platzer, A., Jehl, M.-A., Arnold, R., and Rattei, T. 2015. EffectiveDB— updates and novel features for a better annotation of bacterial secreted proteins and Type III, IV, VI secretion systems. Nucleic Acids Research 44:D669–D674.

Estrada-de Los Santos, P., Palmer, M., Chavez-Ramirez, B., Beukes, C., Steenkamp, E.T., Briscoe, L., Khan, N., Maluk, M., Lafos, M., Humm, E., Arrabit, M., Crook, M., Gross, E., Simon, M.F., Dos Reis Junior, F.B., Whitman, W.B., Shapiro, N., Poole, P.S., Hirsch, A.M., Venter, S.N., and James, E.K. 2018. Whole Genome Analyses Suggests that *Burkholderia* sensu lato Contains Two Additional Novel Genera (*Mycetohabitans* gen. nov., and *Trinickia* gen. nov.): Implications for the Evolution of Diazotrophy and Nodulation in the *Burkholderiaceae*. Genes 9.

Fenselau, S., and Bonas, U. 1995. Sequence and expression analysis of the hrpB pathogenicity operon of *Xanthomonas campestris* pv. *vesicatoria* which encodes eight proteins with similarity to components of the Hrp, Ysc, Spa, and Fli secretion systems. Mol Plant Microbe Interact 8:845–854.

Gsell, M., Fankl, A., Klug, L., Mascher, G., Schmidt, C., Hrastnik, C., Zellnig, G., and Daum, G. 2015. A Yeast Mutant Deleted of GPH1 Bears Defects in Lipid Metabolism. PLOS ONE 10:e0136957.

He, S.Y., Nomura, K., and Whittam, T.S. 2004. Type III protein secretion mechanism in mammalian and plant pathogens. Biochimica et Biophysica Acta 1694:181–206.

Heuer, H., Yin, Y.N., Xue, Q.Y., Smalla, K., and Guo, J.H. 2007. Repeat domain diversity of avrBs3-like genes in *Ralstonia solanacearum* strains and association with host preferences in the field. Appl Environ Microbiol 73:4379–4384.

Hu, Y., Huang, H., Cheng, X., Shu, X., White, A.P., Stavrinides, J., Köster, W., Zhu, G., Zhao, Z., and Wang, Y. 2017. A global survey of bacterial type III secretion systems and their effectors. Environmental Microbiology 19:3879–3895.

Hutin, M., Perez-Quintero, A.L., Lopez, C., and Szurek, B. 2015. MorTAL Kombat: the story of defense against TAL effectors through loss-of-susceptibility. Front Plant Sci 6:535.

Juillerat, A., Bertonati, C., Dubois, G., Guyot, V., Thomas, S., Valton, J., Beurdeley, M., Silva, G.H., Daboussi, F., and Duchateau, P. 2014. BurrH: a new modular DNA binding protein for genome engineering. Scientific Reports 4:3831.

Lackner, G., Moebius, N., and Hertweck, C. 2011a. Endofungal bacterium controls its host by an hrp type III secretion system. The ISME journal 5:252–261.

Lackner, G., Moebius, N., Partida-Martinez, L., and Hertweck, C. 2011b. Complete Genome Sequence of *Burkholderia rhizoxinica*, an Endosymbiont of *Rhizopus microsporus*. Journal of Bacteriology 193:783–784.

Lackner, G., Möbius, N., Scherlach, K., Partida-Martinez, L.P., Winkler, R., Schmitt, I., and Hertweck, C. 2009. Global Distribution and Evolution of a Toxinogenic *Burkholderia-Rhizopus* Symbiosis. Applied and Environmental Microbiology 75:2982–2986.

Ladant, D., and Ullmann, A. 1999. *Bordetella pertussis* adenylate cyclase: a toxin with multiple talents. Trends in Microbiology 7:172–176.

Lastovetsky, O.A., Gaspar, M.L., Mondo, S.J., LaButti, K.M., Sandor, L., Grigoriev, I.V., Henry, S.A., and Pawlowska, T.E. 2016. Lipid metabolic changes in an early divergent fungus govern the establishment of a mutualistic symbiosis with endobacteria. Proceedings of the National Academy of Sciences 113:15102–15107.

Lopez-Fernandez, L., Sanchis, M., Navarro-Rodriguez, P., Nicolas, F.E., Silva-Franco, F., Guarro, J., Garre, V., Navarro-Mendoza, M.I., Perez-Arques, C., and Capilla, J. 2018. Understanding *Mucor circinelloides* pathogenesis by comparative genomics and phenotypical studies. Virulence 9:707–720.

Love, M.I., Huber, W., and Anders, S. 2014. Moderated estimation of fold change and dispersion for RNA-seq data with DESeq2. Genome Biology 15:550.

Mak, A.N.-S., Bradley, P., Cernadas, R.A., Bogdanove, A.J., and Stoddard, B.L. 2012. The Crystal Structure of TAL Effector PthXo1 Bound to Its DNA Target. Science 335:716–719.

Mondo, S.J., Lastovetsky, O.A., Gaspar, M.L., Schwardt, N.H., Barber, C.C., Riley, R., Sun, H., Grigoriev, I.V., and Pawlowska, T.E. 2017. Bacterial endosymbionts influence host sexuality and reveal reproductive genes of early divergent fungi. Nature Communications 8:1843.

Moran, N.A. 1996. Accelerated evolution and Muller’s rachet in endosymbiotic bacteria. Proceedings of the National Academy of Sciences of the United States of America 93:2873–2878.

Moscou, M.J., and Bogdanove, A.J. 2009. A Simple Cipher Governs DNA Recognition by TAL Effectors. Science 326:1501.

Muller, H.J. 1964. THE RELATION OF RECOMBINATION TO MUTATIONAL ADVANCE. Mutation research 106:2–9.

Nazir, R., Mazurier, S., Yang, P., Lemanceau, P., and van Elsas, J.D. 2017. The Ecological Role of Type Three Secretion Systems in the Interaction of Bacteria with Fungi in Soil and Related Habitats Is Diverse and Context-Dependent. Frontiers in Microbiology 8:38.

Partida-Martinez, L.P., and Hertweck, C. 2005. Pathogenic fungus harbours endosymbiotic bacteria for toxin production. Nature 437:884–888.

Partida-Martinez, L.P., Monajembashi, S., Greulich, K.-O., and Hertweck, C. 2007. Endosymbiont-Dependent Host Reproduction Maintains Bacterial-Fungal Mutualism. Current Biology 17:773–777.

Read, A.C., Rinaldi, F.C., Hutin, M., He, Y.-Q., Triplett, L.R., and Bogdanove, A.J. 2016. Suppression of Xo1-Mediated Disease Resistance in Rice by a Truncated, Non-DNA-Binding TAL Effector of *Xanthomonas oryzae*. Frontiers in Plant Science 7.

Römer, P., Recht, S., Strauß, T., Elsaesser, J., Schornack, S., Boch, J., Wang, S., and Lahaye, T. 2010. Promoter elements of rice susceptibility genes are bound and activated by specific TAL effectors from the bacterial blight pathogen, *Xanthomonas oryzae* pv. *oryzae*. New Phytologist 187:1048–1057.

Schandry, N., de Lange, O., Prior, P., and Lahaye, T. 2016. TALE-Like Effectors Are an Ancestral Feature of the *Ralstonia solanacearum* Species Complex and Converge in DNA Targeting Specificity. Frontiers in Plant Science 7.

Schechter, L.M., Roberts, K.A., Jamir, Y., Alfano, J.R., and Collmer, A. 2004. *Pseudomonas syringae* Type III Secretion System Targeting Signals and Novel Effectors Studied with a Cya Translocation Reporter. Journal of Bacteriology 186:543–555.

Schroeder, L., and Ikui, A.E. 2019. Tryptophan confers resistance to SDS-associated cell membrane stress in Saccharomyces cerevisiae. PLoS One 14:e0199484.

South, A. 2011. rworldmap: A New R package for Mapping Global Data. The R Journal 3/1:35–43.

Stella, S., Molina, R., Lopez-Mendez, B., Juillerat, A., Bertonati, C., Daboussi, F., Campos-Olivas, R., Duchateau, P., and Montoya, G. 2014. BuD, a helix-loop-helix DNA-binding domain for genome modification. Acta crystallographica. Section D, Biological crystallography 70:2042–2052.

Szurek, B., Marois, E., Bonas, U., and Van den Ackerveken, G. 2001. Eukaryotic features of the *Xanthomonas* type III effector AvrBs3: protein domains involved in transcriptional activation and the interaction with nuclear import receptors from pepper. The Plant Journal 26:523–534.

Szurek, B., Rossier, O., Hause, G., and Bonas, U. 2002. Type III-dependent translocation of the *Xanthomonas* AvrBs3 protein into the plant cell. Molecular Microbiology 46:13–23.

Van den Ackerveken, G., Marois, E., and Bonas, U. 1996. Recognition of the Bacterial Avirulence Protein AvrBs3 Occurs inside the Host Plant Cell. Cell 87:1307–1316.

