## Supplemental Information for "A TAL effector-like protein of symbiotic *Mycetohabitans* increases stress tolerance and alters the transcriptome of the fungal host *Rhizopus microsporus*"

Materials and Methods
Supplemental Figures
Supplemental Tables
Supplemental References

### Materials and Methods:

#### Bacterial and fungal growth:

*R. microsporus* was grown on half-strength potato dextrose agar (½PDA) at 28°C unless otherwise noted. *Mycetohabitans* spp. were grown at 28°C in liquid or on solid Lysogeny Broth (LB) amended with 1% glycerol and 30 µg/mL ampicillin, with 10 µg/mL gentamycin or 25 µg/mL kanamycin as needed. For cloning purposes, *E. coli* was grown at 37°C on LB amended with 10 µg/mL gentamycin, 25 µg/mL kanamycin, or 10 µg/mL chloramphenicol as needed. Chemically competent *E. coli* cells were used for transformation, while transformation of *Mycetohabitans* spp. was done by electroporation.

To isolate bacteria, one to two-day-old *R. microsporus* cultures were scraped from a plate and ground with a plastic pestle in a microcentrifuge tube. LB was added to the tube and then twice filtered through 0.2 µM filters (Whatman) to remove fungal debris. The filtered liquid was plated with sterile beads and incubated for 5 or more days before colonies appeared. To cure the fungus, *R. microsporus* was successively cultured on liquid and solid media amended with 20 µg/mL ciprofloxacin until no sporulation occurred. Reinfection is carried out by cocultivation of bacterial and fungal partners on solid media as described by Lastovetsky*, et al.* (1) until sporulation is seen.

#### Cloning of *btl* genes and derivatives:

All plasmids used in this study are listed in Table S5, including a series of *btl19-13* Gateway™ entry vectors that were created for use in this study. *btl19-13* was amplified with Q5 High-Fidelity 2X Master Mix (NEB) from *Mycetohabitans* sp. B13 genomic DNA and cloned into SacI-digested pHM1 (2) with NEBuilder HiFi DNA Assembly Master Mix (NEB) to create pSC31 by Gibson Assembly. The entry vector pENTR4 (Thermo Fisher Scientific) and pSC31 were digested with KpnI and EcoRI and the desired bands were gel purified with the Monarch DNA Gel Extraction Kit (NEB), ligated with Quick Ligase (NEB), and transformed into *E. coli* DH5α. The resulting Gateway™ entry vector plasmid, pMEC0, contained *btl19-13* with the native promoter (600 bp), start, and stop codon. Other entry vectors containing derivatives with and without the promoter, start codon, stop codon, or first 45 amino acids were all made with the Q5 Site-Directed Mutagenesis Kit (NEB) and named sequentially as they were created. A plasmid encoding a Btl19-13 derivative with the "RIRK" motif in the C-terminus (aa 692-695) mutated to alanines was similarly created by mutagenesis. For the Btl19-13:mCherry fusion, mCherry was PCR-amplified from pGEM-mCherry with primers that contained SacI restriction enzyme cut sites. The PCR product and pMEC6 were digested with SacI (NEB) and ligated. The resulting pMEC6-mCherry plasmid was transformed into *E. coli* DH5α and encodes Btl19-13:mCherry driven by its native promoter*.* Similar to *btl19-13*, *btl18-14* with a 338 bp promoter was amplified by PCR from *Mycetohabitans sp.* B14 genomic DNA, digested, gel purified, and cloned directly into the entry vector pENTR4 to make pMEC13.

Two expression vectors were modified to be Gateway™ destination vectors for expression in *Mycetohabitans* sp. B13. Gateway™ destination vector cassettes (Thermo Fisher Scientific) were ligated into pBBR1-MCS5 (3) digested with SacI-HF and XhoI and into pCPP6260 (4) digested with SacI-HF and PvuI-HF (NEB) to create pMEF1 and pMEF2, respectively. The resulting plasmids were transformed into *E. coli* DB3.1. LR Clonase II (Thermo Fisher Scientific) was used to recombine pMEC6-mCherry and pMEF2 to create pMEF6, the expression vector for Btl19-13:mCherry. The *btl19-13* complementation plasmid, pBtl19-13, is the product of an LR reaction between pMEC0 and pMEF1. Similarly, pMEC13 was recombined with pMEF1 to make the plasmid pBtl18-14 containing *btl18-14*.

**Microscopy of *R. microsporus* for *btl19-13* expression:**

The plasmid pMEF6 was transformed into *Mycetohabitans* sp. B13 before reinfection into its native *R. microsporus* host. Glycerol stocked spore cultures were suspended in ½ PDB and pipetted onto ½ PDA pads on microscope slides. These were incubated overnight and then stained with DAPI-Fluoromount-G® (SouthernBiotech) the following morning. Slides were imaged on a DeltaVision imaging workstation (Applied Precision). Images were deconvolved with softWoRx software (Applied Precision) and final images were edited with Fiji (5).

**Cya translocation assay and immunoblot:**

Derivatives of *btl19-13* were transferred to pCPP5371 (6) by LR Clonase II (Thermo Fisher Scientific) and transformed into *Pseudomonas syringae* pv. *tomato* (*Pst*) DC3000 or the *Pst* DC3000 *hrcQ-U* mutant (T3SS-) (7). The Cya translocation assay was modified from the protocol described by Chakravarthy, Huot and Kvitko (8). Strains of *Pst* were grown on King's B medium for 2 days at 28°C then suspended in 10 mM $MgCl_2$ to an $OD_{600}$ of 0.5. The youngest fully expanded leaves of 5-week-old *N. benthamiana* plants were infiltrated with a blunt syringe, each strain in triplicate on three separate plants. For each infiltration site, 1 $cm^2$ of infiltrated leaf tissue was harvested 6 hours post infiltration and frozen in liquid nitrogen. Copper BBs (Crosman) were used to grind the tissue by vortexing before 300 µL of 0.1 M HCl was added. Samples were diluted 1:50 to fall within the range of the assay and processed with the Direct cAMP ELISA Kit (Enzo) according to manufacturer's instructions. Total protein content was measured with the Bio-Rad Protein Assay method.

For the corresponding immunoblot, ~1.5 $cm^2$ infiltrated tissue samples were frozen and ground. Protein was extracted in 150 µL of buffer [125 mM Tris pH 8.5, 1% SDS, 10% glycerol] with a fungal protease inhibitor (Millipore Sigma) and resolved by 4-20% Tris Glycine SDS-PAGE (Bio-Rad). Proteins were transferred to a membrane using the Trans-Blot Turbo Transfer System (Bio-Rad). Cya fusion proteins were detected with a 1:300 diluted primary anti-Cya (3D1) mouse monoclonal IgG antibody (Santa Cruz Biotechnology) and a 1:4000 diluted secondary HRP-conjugated goat anti-mouse IgG antibody (Thermo Fisher Scientific). Clarity ECL substrate (Bio-Rad) was used for visualization.

**Microscopy of yeast for Btl19-13 localization:**

Entry vectors with *btl19-13* gene derivatives and previously cloned *Xanthomonas oryzae* pv*. oryzicola* effector Tal1c were recombined into pAG426GAL-eGFP-ccdB (AddGene: #14323; deposited by Susan Lindquist) with LR Clonase II (Thermo Fisher Scientific). For Btl19-13 truncations, the products of this reaction were digested with PspXI and SalI to remove the majority of the Btl19-13 coding region, then self-ligated to contain a little over 2 repeats from the DNA binding domain and the C-terminus (aa 599-714). These constructs were transformed into *Saccharomyces cerevisiae* InvSc1 (Invitrogen) and selected on yeast synthetic uracil drop-out (ura-) media (Sigma). Individual colonies were used to inoculate cultures of liquid ura- media with 2% glucose and incubated overnight at 28°C with shaking. Cells were spun down and resuspended in ura- media containing 0.5 % glucose then allowed to grow for 6 hours. Sterilized 50% galactose was added to the culture to a final concentration of 2% and grown overnight. NucBlue Live ReadyProbes Reagent (Thermo Fisher Scientific) was added to the yeast cells and incubated for at least an hour before imaging by LSM 710 confocal microscope (Zeiss). Final images were edited with Fiji (5).

**Electrophoretic mobility shift assay:**

The coding sequence of Btl19-13 was codon optimized by GenScript for expression in *E. coli* and delivered in pUC57. Btl19-13 was cut from this vector with NotI and BsrGI (NEB) and moved into an expression vector that adds an N-terminal 6xHis-tag to make pFR178. *E. coli*

Rosetta pLysS (DE3) (Novagen) cells were transformed with pFR178. Proteins were expressed and purified as described by Rinaldi, Doyle, Stoddard and Bogdanove (9).

Electrophoretic mobility shift assays were done as previously described using the LightShift Chemiluminescent EMSA Kit (Thermo Scientific) with slight modifications (9). Duplexed oligonucleotides were ordered from Integrated DNA Technologies with biotin labels on the 5' end of the reverse strand. The forward sequences were: binding element (BE) 5'-CGTGGCGTCGTCCACGATTCCGGATTGGACCGACGGCGTC-3' and frame-shifted binding element ($BE^{FS}$) 5'-CGTGGCGTCGTCCACGATTCGGATTGGACCGACGGCGTC-3'. Labeled DNA and purified protein were incubated at room temperature for an hour in binding buffer [10 mM Tris–HCl pH 7.5, 100 mM KCl, 1 mM DTT, 2.5% glycerol, 5 mM $MgCl_2$, 50 ng/μl Poly-dIdC, 0.05% NP-40, 0.4 mM EDTA and 0.1 mg/mL BSA]. Binding reactions were mixed with sample buffer and run on a 1.3% agarose gel at 110 V for ~45 min at 4°C. DNA and DNA-protein complexes were transferred to a positively charged Nylon membrane (Sigma-Aldrich) at 100 V for 30 min at 4°C in 0.5×TBE buffer. The DNA was cross-linked to the nylon membrane by exposure to UV for 12 min in a ChemiDoc imaging system (Bio-Rad). DNA was detected by chemiluminescence following the manufacturer's protocol for the EMSA kit.

**GUS assay:**

A designer TAL effector (dT19-13) was assembled using the Golden Gate TALEN and TAL Effector Kit (AddGene: Kit #1000000024) and additional modules created in the Bogdanove lab for ND, N*, and NS repeats (10). The RVD sequence of dT19-13 was designed to resemble Btl19-13 as closely as possible (Figure S3) with available RVD modules: ND ND NI ND NN NI NG NG N* ND NN NN NS N* NG NN NN NI N*. dT19-13 was assembled into a backbone plasmid (pTAL1) containing the *Xanthomonas oryzae* pv. *oryzicola* effector Tal1c N- and C-termini to create pMEC15. LR Clonase II (Thermo Fisher Scientific) was used to recombine the regions encoding dT19-13 from pMEC15, Btl19-13 from pMEC3, and Tal1c from pCS468 into the binary vector pGWB5, to create pMEC15G, pMEC3G, and pMECtal1cG, respectively. These plasmids were transformed into *Agrobacterium tumefaciens* (*At*) GV3101.

The previously assembled binary vector pCS752 has the pepper Bs3 promoter driving GUS expression with a unique AscI cut site just upstream of the native AvrBs3 binding element (11). The plasmid was digested with AscI (NEB) and dephosphorylated with Antarctic Phosphatase (NEB). Oligonucleotides were ordered from IDT, annealed, phosphorylated, and ligated into the AscI cut site. Two vectors were made: pMEC-GUS1 has the Btl19-13 binding element (BE: 5'-CCACGATTCCGGATTGGAC-3') and pMEC-GUS2 has a scrambled element (sBE: 5'-GTGCACTCGTTAGCCAGAC-3'). Each binding element was preceded by a T residue to be compatible with TAL effectors, though not required for Btl protein binding (12, 13). The constructs were sequenced confirmed for single binding element insertions and transformed into *At* GV3101.

Strains of *At* were used to inoculate liquid cultures and left to grow shaking overnight at 28°C. The next morning, cultures were resuspended in 10 mM $MgCl_2$ with 150 μM of acetosyringone and mixed so each strain was at an $OD_{600}$ of 0.3 in the final mixture. Following a 2-hour incubation, cultures were infiltrated into fully expanded leaves of 5-week-old *N. benthamiana* plants with a needleless syringe after scoring the leaf with a needle. Two leaf punches, each 1 cm in diameter, were harvested 48 hours post infiltration, frozen in liquid nitrogen, and ground with glass beads by vortexing. The ground tissue was suspended in 200 μL of GUS Extraction Buffer [50 mM sodium phosphate (pH 7), 10 mM EDTA, 10 mM β-mercaptoethanol, 0.1% Triton-X100, 0.1% SDS] then vortexed and centrifuged to pellet debris. The supernatant was used in a Bio-Rad Protein Assay to determine total protein concentration and a MUG assay to determine GUS activity.

**Creating a *btl19-13* mutant:**

The 1.6 kb upstream and 1.3 kb downstream flanking regions of *btl19-13* were each amplified by PCR from *Mycetohabitans* sp. B13 genomic DNA and TOPO® cloned into pCR8 (Thermo Fisher Scientific). The construct containing the upstream region was digested with EcoRI-HF and MfeI (NEB), the construct with the downstream region was digested with EcoRI-HF and SalI-HF (NEB), and these were gel purified, ligated, and PCR amplified with OneTaq (NEB) to fuse them before being TOPO® cloned into pCR8. SalI-HF, EcoRI-HF, and XhoI were used to digest the resulting pCR8 vector and EcoRI-HF and SalI-HF was used to digest pK18mobsacB (14), a broad host range vector for use in disrupting genes by sucrose counterselection. The *btl19-13* flanking-region fusion fragment and pK18mobsacB were ligated to create pSC29. *Mycetohabitans* sp. B13 was transformed with pSC29 via electroporation with an overnight recovery period before selection to get single colonies. Transformants were then plated on media amended with 5% sucrose to counterselect for pK18mobsacB loss via double crossover and recover a marker-less mutant, B13Δ*btl19-13*. To create the potentially complemented strains, pBtl19-13 and pBtl18-14 were each transformed into B13Δ*btl19-13*.

***R. microsporus* sporulation assays*:***

Single sporangia from sporulating cultures were plated onto fresh ½ PDA and sealed with micropore paper tape (3M). The plates were kept at room temperature on a laboratory bench. There were 7-10 biological replicates, each from a separate infection event. After 22 days, spores were harvested by scraping the mycelial mat off of the plate surface and vortexing for 2 minutes in 0.00125% Tween-20. Spores were filtered through a sterile cotton swab in a 5 mL serological pipette and quantified twice for two technical replicates using a TC20 Automated Cell Counter (Bio-Rad). Data was analyzed with ANOVA in R (v3.5.3).

For mating plates, the compatible isolates of *R. microsporus*, ATCC 52813 and 52814, were both cured and infected with B13 or B13Δ*btl19-13*. Drops of glycerol stocked spore suspensions were placed on opposite sides of a ½PDA plate and allowed to grow together. Plates were photographed after zygospores started to appear on day 5 or later. Spores were imaged with a Canon EOS attached to an Olympus SZX12.

***R. microsporus* growth assays:**

*R. microsporus* was plated from glycerol stocked spore suspension onto ½ PDA. Two days later, 2 mm x 2 mm patches of mycelia were used to inoculate the center of new plates: water agar, ½ PDA + 500 mM sodium chloride, ½ PDA + 3 mM hydrogen peroxide, or ½ PDA + 0.005% sodium dodecyl sulfate. Each day until growth reached the edge of the plate, colony diameters were measured on 2 axes and the average measurement was recorded. For each repeat, there were 10 plates of fungi infected with each bacterial genotype. The experiment was repeated twice for each condition and set of genotypes. Data from each repeat was analyzed with ANOVA in R and each repeat showed the same trend.

**RNAseq:**

Glycerol stocked spore suspensions were plated on to ½ PDA and incubated for 24 hours. Half of the vegetative growth from 3 plates was pooled for each biological replicate. Three biological replicates were done for each of the three bacterial genotypes. RNA extractions were done with RNeasy Plant Mini Kit (Qiagen) according to manufacturer's protocol, except for sample homogenization. Tissue was directly placed into a 1.5 mL tube containing 450 uL of RLT buffer (subsidized with DTT as per kit instructions) and two Copper BBs (Crosman). Tubes were vortexed for 10 sec before incubating at 55°C for 5 min, then vortexted again. This slurry was used as the input for the kit. RNA quality and quantity was assessed by agarose gel, NanoDrop (Thermo Fisher Scientific), and Qubit Fluorometer (Invitrogen) before being sent to Novogene for eukaryotic library preparation and paired-end (2 x 150 bp) Illumina sequencing. Each sample library yielded >20 million reads (>6 Gb). RNA-seq analysis was carried out with Trimmomatic for quality control and trimming adapter sequences followed by HISAT2 (15) to align the reads to the *R. microsporus* ATCC 52813 genome assembly Rhimi1_1 (16). StringTie (17) was used to generate gene counts which were fed into DESeq2 (18) for expression analysis. Genes were considered differentially expressed for an adjusted $p$ value $<0.05$ and a log2foldchange$>|1|$. Binding element (BE) prediction was done with TALE-NT 2.0 on the 500 bp promoter regions of the differentially expressed genes, with the option of a T or C in the -1 position (19).

**Genomic DNA preparation and Southern blots:**

Three milliliters of turbid liquid cultures of bacteria were used in the MasterPure™ Gram Positive DNA Purification Kit (Lucigen) according to manufacturer's instructions. For strains that were newly isolated, we tested to see if they were *Mycetohabitans spp*. by PCR using previously developed 23S primers that are specific for *Burkholderia* and related species: GlomGiGF and LSU483r (20). PCR fragments were then gel-purified and sequenced to confirm. When we could not successfully extract bacteria from a fungal isolate with no known symbiont, we extracted fungal genomic DNA using the E.Z.N.A. Fungal Mini Kit (Omega) and attempted the same PCR to verify that no bacteria were present.

Southern blot analysis was done using the Amersham ECL Direct Labeling and Detection Kit (GE Healthcare Life Sciences). About 2.5 µg of genomic DNA was digested overnight with the restriction enzyme AatII (NEB), then run for 18 hours at 35 V on a 1% agarose gel at 4˚C. The gel was post-stained in a 3X GelRed solution to visualize equal loading and digest efficiency. The DNA was depurinated with a 10-minute wash in 0.25 M HCl and washed in denaturation buffer [250 µM NaOH, 1.5 mM NaCl] before being transferred to an Amersham Hybond N+ nylon membrane Kit (GE Healthcare Life Sciences) overnight. The blot was crosslinked and then hybridized at 60˚C overnight with a 2.1 kB probe of *btl19-13* amplified from pMEC0. The membrane was washed twice for ten minutes at 60˚C in Primary Wash Buffer [2 M Urea, 0.1% SDS, 50 mM Sodium Phosphate pH 7, 150 mM NaCl, 0.1 mM $MgCl_2$, 2% Blocking Reagent supplied with kit] and twice for five minutes in Secondary Wash Buffer [1 M Tris base, 2 M NaCl]. It was then incubated with Amersham CDP Star Detection Reagent Kit (GE Healthcare Life Sciences) for 4 minutes and visualized on a ChemiDoc imaging system (Bio-Rad) under signal accumulation mode with the hi-resolution blot sensing program.

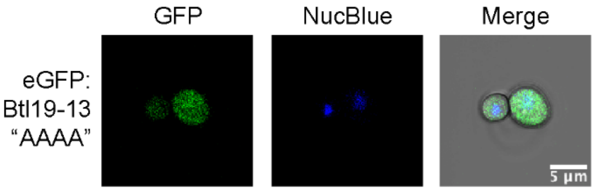


**Figure S1. Expression of full-length Btl19-13 with an NLS mutation in yeast cells.** Confocal microscopy of *Saccharomyces cerevisiae* expressing eGFP:Btl19-13 (“RIRK” to “AAAA”; aa 692-695) and stained with NucBlue to indicate nuclei. Protein expression was induced with galactose 24 hours prior to imaging.

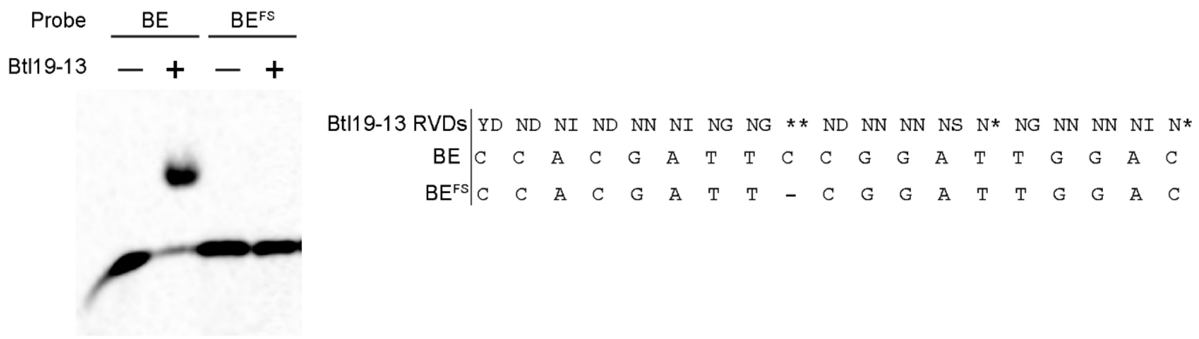


**Figure S2. Binding of Btl19-13 to two predicted binding elements**. Electrophoretic mobility shift assay of Btl19-13 with BE, the binding element that is predicted to be the best following the TAL effector RVD code, or $BE^{FS}$, the same binding element with a frameshift where the ** RVD would bind. For each binding reaction, 150 pM of DNA probe was added. The addition of 100 nM protein is indicated with a plus (+) or minus (-). To the right are the sequences of BE and $BE^{FS}$ as they correspond to the RVDs of Btl19-13. An asterisk (*) indicates a missing amino acid in the RVD.

| | | | | | | | | | | | | | | | | | | | |
|---|---|---|---|---|---|---|---|---|---|---|---|---|---|---|---|---|---|---|---|
| Btl19-13 RVDs | YD | ND | NI | ND | NN | NI | NG | NG | ** | ND | NN | NN | NS | N* | NG | NN | NN | NI | N* |
| BE | C | C | A | C | G | A | T | T | C | C | G | G | A | T | T | G | G | A | C |
| dT19-13 RVDs | ND | ND | NI | ND | NN | NI | NG | NG | N* | ND | NN | NN | NS | N* | NG | NN | NN | NI | N* |
| BE | C | C | A | C | G | A | T | T | C | C | G | G | A | T | T | G | G | A | C |
| sBE | G | T | G | C | A | C | T | C | G | T | T | A | G | C | C | A | G | A | C |

**Figure S3. Alignment of Btl19-13 and dT19-13 RVDs with the binding element (BE) and scrambled binding element (sBE) used in the GUS assay.** An asterisk (*) indicates a missing amino acid in the RVD based on alignment with other repeats.

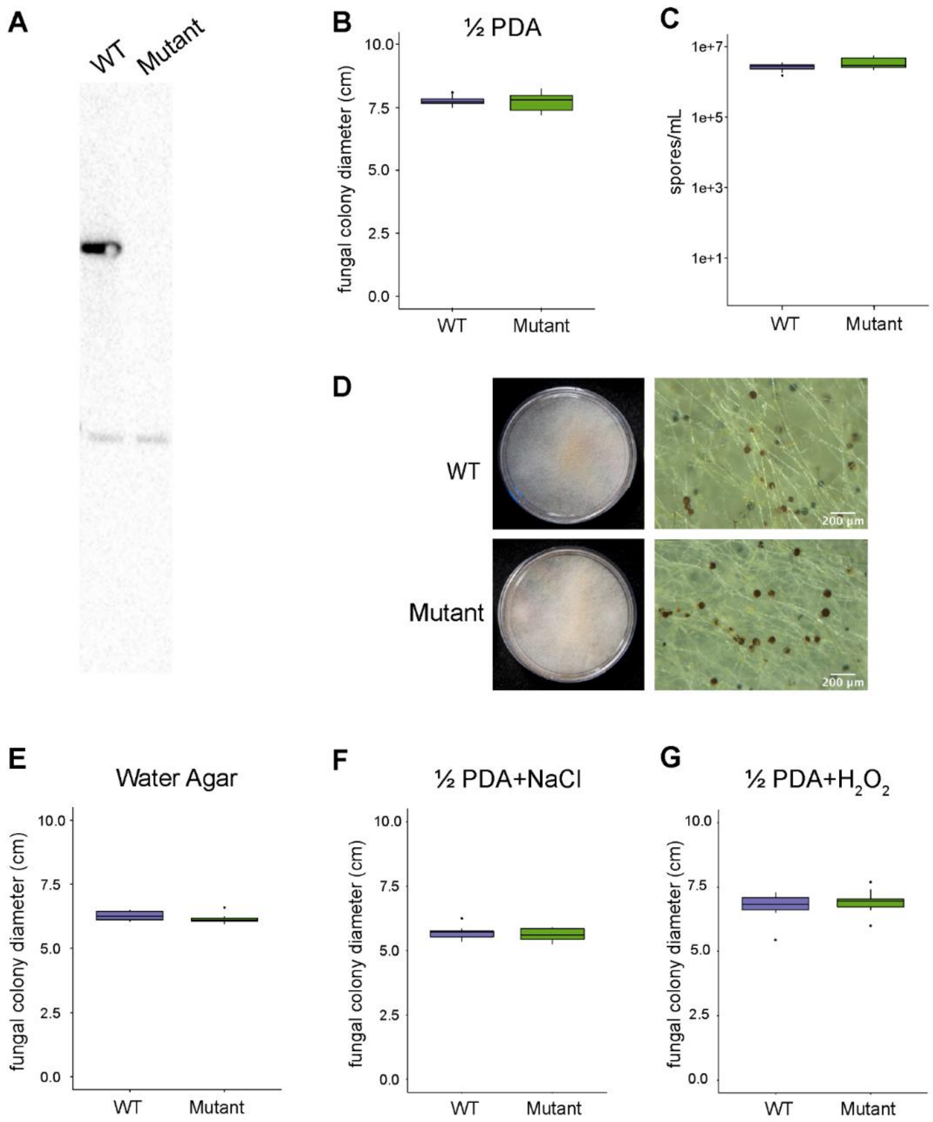


**Figure S4. Sporulation and growth of fungi infected with B13 and a *btl19-13* mutant. (A)** Southern blot of genomic DNA confirming that *btl19-13* is not in the mutant strain B13Δ*btl19-13*, though the separate *btl* gene fragment remains*.* **(B)** Growth of *R. microsporus* on half-strength potato dextrose agar (½ PDA), after four days of growth. Data represents 7-10 biological replicates, each started with a sporangium from separate infection plates. **(C)** Spores were counted from the same plates in (B) after 21 days. **(D)** Photo of sporulating mating plates. A spore suspension of *R. microsporus* ATCC 52814 and ATCC 52813 infected with B13 or B13Δ*btl19-13* was each placed on one side of the plate, and they were allowed to grow together for 5 days at 28°C. **(E-G)** Fungal colony diameter of *R. microsporus* after growing for 3 days on water-agar, 5 days on ½ PDA with 500 mM sodium chloride, and 3 days on ½ PDA with 3 mM hydrogen peroxide at 28°C. Data represent one repeat of 10 biological replicates, each started with a 2 mm x 2 mm plug from a two-day old plate started from glycerol stocked spores. These experiments were repeated twice with similar results. All data were analyzed by ANOVA with a post-hoc Tukey's test ($p < 0.05$).

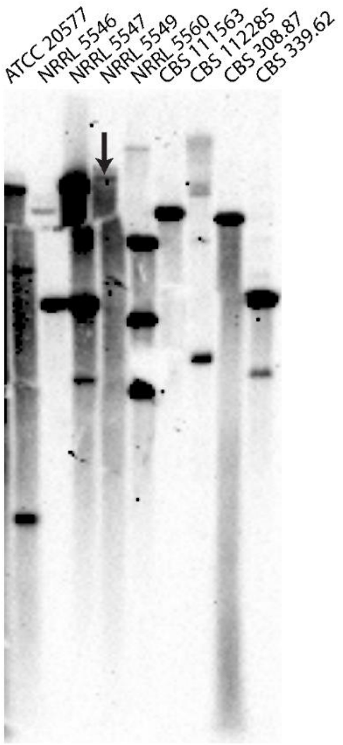


**Figure S5. Increased exposure time of a Southern blot from Figure 4B.** Southern blot of genomic DNA prepared from 9 *Mycetohabitans* spp., probed with *btl19-13* amplified from B13. Strains are identified by the culture collection accession of their fungal hosts. An arrow indicates the band that was not clearly visible in the shorter exposure in Figure 4B.

**Table S1. Information about *btl* genes from 3 genomes of *Mycetohabitans* spp.**

| Btl Protein | Repeats | Strain | Gene Name | Plasmid or Chromosome | HrpB Box (bp upstream) | NLS | Repeat Variable Diresidue (RVD) Sequence |
|---|---|---|---|---|---|---|---|
| **Btl28_1** | 28 | B1 | RBRH_01770 | Plasmid 2 (173 kb) | -239 to -216 | RIRK | YD NG ** NN NG ** NN NI NS NI NS NI NN NG ** NK NA NI NS NI NI NI ND NS NG KG NT NG |
| **Btl21_1** | 21 | B1 | RBRH_01844 | Plasmid 1 (822 kb) | -230 to -207 | RIRK | YD NI NI NN NT NN NI NI NN ND NI NT NR NS ND NN NG NG NS N* |
| **Btl7_1** | 7 | B1 | RBRH_01778 | Plasmid 2 (173 kb) | -219 to -196 | QIRK | HD NG NI NG NG NG NN |
| **Btl19_13** | 19 | B13 | SAMN04487768_ 2744 | Chromosome | -239 to -216 | RIRK | YD ND NI ND NN NI NG NG ** ND NN NN NS N* NG NN NN NI N* |
| **Btl18_14** | 18 | B14 | SAMN04487769_ 0211 | Plasmid | -239 to -216 | RIRK | YD NG NI NI NG ** ND ND NI NS NN NS N* NG NN NN NI N* |
| **Btl15_14c** | 15 | B14 | SAMN04487769_ 1929 | Chromosome | -239 to -216 | RIRK | YD ND NI NI NG ** ND NN NN NS N* NG NN NI N* |
| **Btl15_14p** | 15 | B14 | SAMN04487769_ 0204 | Plasmid | -219 to -196 | RIRK | YD NG NG NG NG NI NI NI NI NI NI S* NN NN NI |

**Table S2. Predicted promoter regulatory elements for *btl* genes.**

| BTL | Predicted HrpB Box | Predicted HrpB Box Location (bp upstream of start codon) | Predicted -10 Element |
|---|---|---|---|
| **Btl28_1** | TTCGctcagtaggtgaagcgTTCG | -239 to -216 | caaagt |
| **Btl21_1** | TTCGttcagcacgcaatgcgTTCG | -230 to -207 | caaagt |
| **Btl7_1** | TTCGccggctcggcggtgcgTTCG | -219 to -196 | caaggt |
| **Btl19_13** | TTCGcccagtaggtgaagtgTTCG | -239 to -216 | caaagt |
| **Btl18_14** | TTCGcccagtaggtgaagtgTTCG | -239 to -216 | caaagt |
| **Btl15_14c** | TTCGcccagtaggtgaagcgTTCG | -239 to -216 | caaagt |
| **Btl15_14p** | TTCGccggatcggtggcgcgTTCG | -219 to -196 | caaggt |

**Table S3. Genes that were differentially expressed in *R. microsporus* infected with wildtype B13 (WT) and complement, B13Δ*btl19-13*(pBtl19-13) (Comp), compared to the mutant, B13Δ*btl19-13* (Mut).**

| Gene | Gene Description | log2FC WT vs Mut | log2FC Comp Vs Mut | BE Ratio Prediction[a] | BE Start Position |
|---|---|---|---|---|---|
| **RHIMIDRAFT_298141** | *hypothetical protein* | 7.69 | 7.96 | 2.31 | 219 |
| **RHIMIDRAFT_253996** | *hypothetical protein* | 5.94 | 6.56 | 2.82 | 275 |
| **RHIMIDRAFT_266868** | *hypothetical protein* | 5.52 | 6.37 | - | - |
| **RHIMIDRAFT_232418** | *DUF1960-domain-containing protein* | 3.76 | 5.23 | 2.97 | 91 |
| **RHIMIDRAFT_313464** | *hypothetical protein* | 2.85 | 3.05 | 2.92 | 218 |
| **RHIMIDRAFT_267780** | *hypothetical protein* | 2.05 | 2.07 | 2.89 | 88 |
| **RHIMIDRAFT_283293** | *hypothetical protein* | 1.65 | 1.92 | 2.71 | 172 |
| **RHIMIDRAFT_206793** | *4-dihydromethyltrisporate dehydrogenase* | 2.30 | 1.79 | 2.87 | 297 |
| **RHIMIDRAFT_208941** | *DNA clamp loader* | 1.06 | 1.27 | 2.44 | 143 |
| **RHIMIDRAFT_275463** | *hypothetical protein* | -1.32 | -1.06 | 2.25 | 324 |
| **RHIMIDRAFT_119095** | *hypothetical protein* | -1.06 | -1.33 | 2.75 | 37 |
| **RHIMIDRAFT_279123** | *hypothetical protein* | -2.22 | -1.62 | 2.70 | 203 |
| **RHIMIDRAFT_232860** | *hypothetical protein* | -3.60 | -3.11 | 2.88 | 297 |
| **RHIMIDRAFT_264374** | *hypothetical protein* | -5.40 | -3.48 | 2.69 | 218 |
| **RHIMIDRAFT_274095** | *hypothetical protein* | -5.05 | -3.89 | 2.32 | 424 |

[a] Best binding element (BE) for BTL19-13 predicted in the 500 bp promoter region of this gene by TALE-NT (19); lower scores are better.

**Table S4. Isolates of *Rhizopus microsporus* and *Mycetohabitans* spp. used or identified in this study.**

| Accession | Taxon | Origin | Bacterial Species | Strain Name | Bacterial Presence Determined |
|---|---|---|---|---|---|
| ATCC 62417 | *Rhizopus microsporus* van Tieghem | Rice seedlings, Japan | *Mycetohabitans rhizoxinica* | HKI 0454/B1 | Partida-Martinez and Hertweck (21) |
| ATCC 20577 | *Rhizopus sp.* strain F-1360 | Soil, Japan | *Mycetohabitans sp.* | HKI 0512/B2 | Lackner, *et al.* (22) |
| NRRL 13479; CBS 699.68; ATCC 52813 | *Rhizopus microsporus* van Tieghem | Soil, Ukraine | *Mycetohabitans endofungorum* | HKI 0402/ B4/B13 | Lackner, *et al.* (22) Dolatabadi, *et al.* (23); |
| NRRL 13478; CBS 700.68; ATCC 52814 | *Rhizopus microsporus* van Tieghem | Forest soil, Georgia | *Mycetohabitans endofungorum* | HKI 0403/ B7/B14 | Lackner, *et al.* (22) Dolatabadi, *et al.* (23); |
| CBS 261.28; ATCC 52811 | *Rhizopus microsporus* van Tieghem | Unspecified, USA | *Mycetohabitans rhizoxinica* | HKI 0513/B6 | Lackner, *et al.* (22) Dolatabadi, *et al.* (23); |
| CBS 262.28; ATCC 52812 | *Rhizopus microsporus* var Chinensis | Unspecified, USA | *Mycetohabitans sp.* | - | This study |
| NRRL 5546 | *Rhizopus microsporus* | Soil, Brazil | *Mycetohabitans sp.* | - | This study |
| NRRL 5547 | *Rhizopus microsporus* | Soil, Philippines | *Mycetohabitans sp.* | - | This study |
| NRRL 5549; ATCC 46352 | *Rhizopus microsporus* | Rabbit Dung, Wisconsin, USA | *Mycetohabitans sp.* | - | This study |
| NRRL 5560 | *Rhizopus microsporus* | Corn, USA | *Mycetohabitans sp.* | - | This study |
| CBS 112285 | *Rhizopus microsporus* Tieghem | Ground nuts, Mozambique | *Mycetohabitans endofungorum* | HKI 0456/B5 | Lackner, *et al.* (22) |
| CBS 111563 | *Rhizopus microsporus* Tieghem | Sufu starter culture, Vietnam | *Mycetohabitans endofungorum* | HKI 0455/B3 | Lackner, *et al.* (22) Dolatabadi, *et al.* (23); |
| CBS 339.61 | *Rhizopus microsporus* var. *oligosporus* (Saito) Schipper & Stalpers | Tempeh, Indonesia | *Mycetohabitans rhizoxinica* | - | Dolatabadi, *et al.* (23); |
| NRRL 28628; CBS 308.87 | *Rhizopus microsporus* van Tieghem | Necrotic tissue from a man’s hand, Australia | *Mycetohabitans sp.* | HKI 0404/B8 | Lackner, *et al.* (22) |

**Table S4 cont. Isolates of *Rhizopus microsporus* and *Mycetohabitans* spp. used or identified in this study.**

| Accession | Taxon | Origin | Bacterial Species | Strain Name | Bacterial Presence Determined |
|---|---|---|---|---|---|
| CBS 113206 | *Rhizopus microsporus* var. *tuberosus* | Koji beer, China | None | NA | This study |
| NRRL 1508 | *Rhizopus microsporus* | Soil, USA | None | NA | This study |

**Table S5. Plasmids used in this study with a description of their relevant features and their source.**

| PLASMID | DESCRIPTION | SOURCE |
|---|---|---|
| **pK18mobsacB** | Broad host range marker-less deletion vector | Schafer*, et al.* (14) |
| **pSC29** | pK18mobsacB with *btl19-13* flanking regions | This study |
| **pHM1** | Broad host range cosmid vector | Hopkins, White, Choi, Guo and Leach (2) |
| **pSC31** | pHM1 containing *btl19-13* with native promoter | This study |
| **pMEC0** | pENTR4-Btl19-13 with native promoter, native start codon, and native stop codon in pENTR4 (Invitrogen) | This study |
| **pMEC1** | pENTR4-Btl19-13 without native promoter or stop codon, but with start codon | This study |
| **pMEC2** | pENTR4-Btl19-13 without native promoter or start codon, but with stop codon | This study |
| **pMEC5** | pMEC2 with RIRK NLS mutated to AAAA (aa 692-695) | This study |
| **pMEC6** | pMEC0 with no stop codon; *btl19-13* with native promoter and native start codon | This study |
| **pMEC6-mCherry** | *btl19-13* with native promoter, native start codon, and a C-terminal mCherry fusion | This study |
| **pMEC7** | pMEC3 with N-terminal 45 aa truncation | This study |
| **pMEC8** | pMEC3 with only first 45 aa | This study |
| **pMEC13** | *btl18-14* with native promoter, native start codon, and native stop codon | This study |
| **pBBR1-MCS5** | Broad host range expression vector with multiple cloning site | Obranić, Babić and Maravić-Vlahoviček (3) |
| **pMEF1** | pBBR1-MCS5 with a Gateway Destination vector cassette added at the MCS | This study |
| **pBtl19-13** | pMEF1 containing *btl19-13* with native promoter from pMEC0 | This study |
| **pBtl18-14** | pMEF1 containing *btl18-14* with native promoter from pMEC13 | This study |
| **pCPP6260** | NptII promoter driving EYFP expression | Worley (4) |
| **pMEF2** | pCPP6260 with a Gateway Destination vector cassette | This study |
| **pMEF6** | pMEF2 encoding Btl19:13:mCherry with its native promoter | This study |
| **pCPP5371** | Gateway destination vector for Cya fusion proteins | Oh, Kvitko, Morello and Collmer (6) |
| **pCPP5388** | pCPP5371 expressing AvrPto:Cya | Lam*, et al.* (24) |
| **pMEC1C** | pCPP5371 expressing Btl19-13:Cya | This study |
| **pMEC7C** | pCPP5371 expressing Btl19-13[45:]:Cya | This study |
| **pMEC8C** | pCPP5371 expressing Btl19-13[1:45]:Cya | This study |
| **pAG426GAL-EGFP-ccdb** | Gateway destination vector for yeast expression; GAL1 promoter driving N-terminal eGFP tagged expression | Addgene plasmid # 14323 deposited by Susan Lindquist |
| **pCS468** | A Gateway entry vector containing Tal1c cloned from *Xanthomonas oryzae* pv. *oryzicola* | C. Schmidt |
| **pYeGFP-TAL1C** | pAG426GAL-eGFP-tal1c | This study |
| **pMEC2Y** | pAG426GAL-eGFP-Btl19-13 | This study |
| **pMEC5Y** | pAG426GAL-eGFP-Btl19-13 NLS mutant; RIRK mutated to AAAA (aa 692-695) | This study |

**Table S5** cont.

| PLASMID | DESCRIPTION | SOURCE |
|---|---|---|
| **pMEC2YC** | pAG426GAL-eGFP-Btl19-13[599-714] | This study |
| **pMEC5YC** | pAG426GAL-eGFP-Btl19-13[599-714] NLS mutant; RIRK mutated to AAAA (aa 692-695) | This study |
| **pFR178** | Expression of codon-optimized Btl19-13 with a N-terminal 6xHis tag under lac induction | This study |
| **pTAL1** | A gateway entry vector with the Tal1c N- and C-termini; created as part of the kit to make designer TAL effectors | Cermak, *et al.* (10) |
| **pMEC15** | Gateway entry vector with dT19-13; pTAL1 with the RVD sequence most closely related Btl19-13 assembled in between the Tal1c N- and C- termini | This study |
| **pGWB5** | Binary Gateway Destination vector with a 35S promoter | Nakagawa, *et al.* (25) |
| **pCS752** | Binary vector with 343 bp Bs3 promoter from pepper driving GUS expression | Doyle, *et al.* (11) |
| **pGWB5-AvrBs3** | pGWB5 expressing AvrBs3 cloned from *Xanthomonas euvesicatoria* | Cernadas, *et al.* (26) |
| **pMEC3G** | pGWB5 expressing Btl19-13 from pMEC3 | This study |
| **pMEC15G** | pGWB5 expressing dT19-13 from pMEC15 | This study |
| **pMECtal1cG** | pGWB5 expressing Tal1c from pCS468 | This study |
| **pMEC-GUS1** | pCS752 with a Btl19-13 binding element inserted into the AscI cut site | This study |
| **pMEC-GUS2** | pCS752 with a scrambled binding element inserted into the AscI cut site | This study |

### Supplemental References


1. Lastovetsky OA*, et al.* (2016) Lipid metabolic changes in an early divergent fungus govern the establishment of a mutualistic symbiosis with endobacteria. *Proceedings of the National Academy of Sciences* 113(52):15102-15107.
2. Hopkins CM, White FF, Choi SH, Guo A, & Leach JE (1992) Identification of a family of avirulence genes from *Xanthomonas oryzae* pv. *oryzae*. *Mol Plant Microbe Interact* 5(6):451-459.
3. Obranić S, Babić F, & Maravić-Vlahoviček G (2013) Improvement of pBBR1MCS plasmids, a very useful series of broad-host-range cloning vectors. *Plasmid* 70(2):263-267.
4. Worley J (2013) Investigation Of The Interconnected Roles Of CmaL And HopAA1-1 In The Virulence Of *Pseudomonas Syringae* pv.*tomato* DC3000 In *Nicotiana Benthamiana*. Ph. D. (Cornell University, Ithaca, NY).
5. Schindelin J*, et al.* (2012) Fiji: an open-source platform for biological-image analysis. *Nature Methods* 9(7):676-682.
6. Oh H-S, Kvitko BH, Morello JE, & Collmer A (2007) *Pseudomonas syringae* Lytic Transglycosylases Coregulated with the Type III Secretion System Contribute to the Translocation of Effector Proteins into Plant Cells. *Journal of Bacteriology* 189(22):8277-8289.
7. Badel JL, Shimizu R, Oh HS, & Collmer A (2006) A *Pseudomonas syringae* pv. *tomato* avrE1/hopM1 mutant is severely reduced in growth and lesion formation in tomato. *Mol Plant Microbe Interact* 19(2):99-111.
8. Chakravarthy S, Huot B, & Kvitko BH (2017) Effector Translocation: Cya Reporter Assay. *Bacterial Protein Secretion Systems: Methods and Protocols*, eds Journet L & Cascales E (Springer New York, New York, NY), pp 473-487.
9. Rinaldi FC, Doyle LA, Stoddard BL, & Bogdanove AJ (2017) The effect of increasing numbers of repeats on TAL effector DNA binding specificity. *Nucleic Acids Research* 45(11):6960-6970.
10. Cermak T*, et al.* (2011) Efficient design and assembly of custom TALEN and other TAL effector-based constructs for DNA targeting. *Nucleic Acids Research* 39(12):e82.
11. Doyle EL*, et al.* (2013) TAL effector specificity for base 0 of the DNA target is altered in a complex, effector- and assay-dependent manner by substitutions for the tryptophan in cryptic repeat -1. *PloS one* 8(12):e82120-e82120.
12. Boch J*, et al.* (2009) Breaking the code of DNA binding specificity of TAL-type III effectors. *Science* 326(5959):1509-1512.
13. de Lange O*, et al.* (2014) Programmable DNA-binding proteins from *Burkholderia* provide a fresh perspective on the TALE-like repeat domain. *Nucleic Acids Res* 42(11):7436-7449.
14. Schafer A*, et al.* (1994) Small mobilizable multi-purpose cloning vectors derived from the *Escherichia coli* plasmids pK18 and pK19: selection of defined deletions in the chromosome of *Corynebacterium glutamicum*. *Gene* 145(1):69-73.
15. Kim D, Paggi JM, Park C, Bennett C, & Salzberg SL (2019) Graph-based genome alignment and genotyping with HISAT2 and HISAT-genotype. *Nature Biotechnology* 37(8):907-915.
16. Mondo SJ*, et al.* (2017) Bacterial endosymbionts influence host sexuality and reveal reproductive genes of early divergent fungi. *Nature Communications* 8(1):1843.

17. Pertea M, Kim D, Pertea GM, Leek JT, & Salzberg SL (2016) Transcript-level expression analysis of RNA-seq experiments with HISAT, StringTie and Ballgown. *Nature Protocols* 11(9):1650-1667.
18. Love MI, Huber W, & Anders S (2014) Moderated estimation of fold change and dispersion for RNA-seq data with DESeq2. *Genome Biology* 15(12):550.
19. Doyle EL*, et al.* (2012) TAL Effector-Nucleotide Targeter (TALE-NT) 2.0: tools for TAL effector design and target prediction. *Nucleic Acids Research* 40(W1):W117-W122.
20. Mondo SJ, Toomer KH, Morton JB, Lekberg Y, & Pawlowska TE (2012) Evolutionary stability in a 400-million-year-old heritable facultative mutualism. *Evolution* 66(8):2564-2576.
21. Partida-Martinez LP & Hertweck C (2005) Pathogenic fungus harbours endosymbiotic bacteria for toxin production. *Nature* 437(7060):884-888.
22. Lackner G*, et al.* (2009) Global Distribution and Evolution of a Toxinogenic *Burkholderia-Rhizopus* Symbiosis. *Applied and Environmental Microbiology* 75(9):2982-2986.
23. Dolatabadi S*, et al.* (2016) Food preparation with mucoralean fungi: A potential biosafety issue? *Fungal Biology* 120(3):393-401.
24. Lam HN*, et al.* (2014) Global Analysis of the HrpL Regulon in the Plant Pathogen *Pseudomonas syringae* pv. *tomato* DC3000 Reveals New Regulon Members with Diverse Functions. *PLoS ONE* 9(8):e106115.
25. Nakagawa T*, et al.* (2007) Development of series of gateway binary vectors, pGWBs, for realizing efficient construction of fusion genes for plant transformation. *Journal of Bioscience and Bioengineering* 104(1):34-41.
26. Cernadas RA*, et al.* (2014) Code-Assisted Discovery of TAL Effector Targets in Bacterial Leaf Streak of Rice Reveals Contrast with Bacterial Blight and a Novel Susceptibility Gene. *PLOS Pathogens* 10(2):e1003972.
